# Three dimensional fibrotic extracellular matrix directs microenvironment fiber remodeling by fibroblasts

**DOI:** 10.1101/2023.08.09.552411

**Authors:** Mehmet Nizamoglu, Frederique Alleblas, Taco Koster, Theo Borghuis, Judith M. Vonk, Matthew J. Thomas, Eric S. White, Carolin K. Watson, Wim Timens, Karim C. El Kasmi, Barbro N. Melgert, Irene H. Heijink, Janette K. Burgess

**Affiliations:** University of Groningen, University Medical Center Groningen, Department of Pathology and Medical Biology, Groningen, the Netherlands; University of Groningen, University Medical Center Groningen, Groningen Research Institute for Asthma and COPD (GRIAC), Groningen, the Netherlands; University of Groningen, University Medical Center Groningen, Department of Epidemiology, Groningen, the Netherlands; Immunology & Respiratory Diseases Research, Boehringer Ingelheim Pharma GmbH & Co. KG, Biberach an der Riss, Germany; Boehringer Ingelheim Pharmaceuticals, Inc., Ridgefield, CT, United States; University of Groningen, Department of Molecular Pharmacology, Groningen Research Institute for Pharmacy, Groningen, the Netherlands; University of Groningen, University Medical Center Groningen, Department of Pulmonology, Groningen, the Netherlands; University of Groningen, University Medical Center Groningen, W.J. Kolff Institute for Biomedical Engineering and Materials Science-FB41, Groningen, the Netherlands

**Keywords:** fibrosis, ECM-derived hydrogels, collagen organization, crosslinking, biomechanics, idiopathic pulmonary fibrosis

## Abstract

Idiopathic pulmonary fibrosis (IPF), for which effective treatments are limited, results in excessive and disorganized deposition of an aberrant extracellular matrix (ECM). An altered ECM microenvironment is postulated to contribute to disease perpetuation in a feed-forward manner through inducing profibrotic behavior by lung fibroblasts, the main producers and regulators of ECM. Here, we examined this hypothesis in a 3D *in vitro* model system by growing primary human lung fibroblasts in ECM-derived hydrogels from non-fibrotic (control) or IPF lung tissue. Culture of fibroblasts in fibrotic hydrogels did not trigger a change in the overall amount of collagen or glycosaminoglycans but did cause a drastic change in fiber organization compared to culture in control hydrogels. Mechanical properties of fibrotic hydrogels were modified by fibroblasts while control hydrogels were not. These results illustrate how the 3D microenvironment plays a crucial role in directing cells to exhibit pro-fibrotic responses by providing biochemical and/or biomechanical cues.

## Introduction

Tissue fibrosis results from an increase in fibroblasts with an aberrant deposition of extracellular matrix (ECM) and abnormal alterations of the ECM structure and composition [1]. While fibrosis is recognized as a coinciding phenomenon in some diseases, such as in inflammatory diseases or several cancers, organ fibrosis itself is one of the leading causes of death worldwide each year [2]. Among these diseases, idiopathic pulmonary fibrosis (IPF) has a worse prognosis than most cancers and remains incurable to date [3]. Currently, IPF is thought to originate from repeated (micro)injuries to the lung epithelium resulting in an aberrant tissue repair response [4]. During this anomalous tissue repair response, fibroblasts emerge as key players that deposit ECM in an abnormal manner, resulting in scarring of lung interstitium that impairs gas exchange in lungs of patients with IPF [5]. Although there is an urgent unmet need for developing novel treatment strategies, lack of appropriate animal models that recapitulate this human disease hinders this process [6]. For new and improved therapeutics targeting IPF, our understanding of how fibrotic responses are perpetuated and how IPF progresses needs to be advanced.

ECM is drastically altered in fibrotic lung diseases both biochemically and biomechanically [7]. While collagen deposition in the alveolar septa is considered one of the hallmarks of fibrotic scar development, numerous other ECM components such as fibronectin, hyaluronic acid, periostin and fibulin-1 are also present to a greater extent in fibrotic lung ECM [8]. In addition to altered ECM composition, fiber structure in fibrotic ECM is also substantially different compared to healthy ECM: fibrotic lungs having a higher percentage of disorganized collagen [9, 10]. Such changes in the fiber organization and content are also postulated to translate into the well-documented changes in the mechanical properties of fibrotic tissue: IPF lungs are many-fold stiffer than control counterparts [11]. Recently, decreased stress relaxation properties of fibrotic lungs were also described, illustrating not only stiffness but also additional mechanical parameters accompany lung fibrosis [12]. Although initially thought of as an inert structure that only provided a physical scaffold, ECM has now been shown to instruct behavior of resident and transmigratory cells [13]. ECM deposited by fibroblasts in fibrosis resulted in activation of naïve fibroblasts seeded onto this ECM [14]. In addition to the origin of the microenvironment, the dimensionality of the environment (two-dimensional (2D) vs. three-dimensional (3D)) has been shown to influence how fibroblasts respond to their microenvironment [15]. While these pioneering studies illustrate that a fibrotic microenvironment instructs cellular behavior, the influence of a 3D fibrotic microenvironment on fibroblasts remains less explored.

Hydrogels, which are water-swollen polymeric networks, have been used as an *in vitro* tool to mimic the 3D organ microenvironment. Synthetic hydrogels, such as those based on dextran [15], and natural hydrogels based on collagen type I [9] have been used for *in vitro* studies. While synthetic hydrogels provide opportunities to fine-tune the structural arrangement of the fibers and mechanical properties, they lack the biological implications of the altered biochemical microenvironment as found in IPF lungs [16]. Natural hydrogels can provide bioactive cues to cells encapsulated in them; however, mechanically tuning these hydrogels to mimic a diseased microenvironment is rather limited [17]. Hydrogels made of decellularized organ-derived ECM can provide an ideal background for addressing these concerns [18-20]. Decellularized ECMs (dECMs) retain most of the biochemical composition of the native organs and tissues [21], and hydrogels derived from these dECMs have been shown to recapitulate the mechanical properties of their native tissues [18, 19]. Specifically, ECM-derived hydrogels prepared from lung tissue from patients with IPF show the increased stiffness characteristics of native tissue, making these hydrogels an ideal candidate for recapitulating the (fibrotic) microenvironment *in vitro* [12]. Previous studies utilizing poly(ethylene glycol) (PEG)-hybrid murine [22], porcine [23] or human [24] lung ECM-derived hydrogels illustrated ECM-fibroblast dynamics in 2D cultures by comparing the responses of fibroblasts seeded onto these native (soft) and modified (stiff) ECM-derived hydrogels. The unexplored interaction between the 3D fibrotic ECM and fibroblasts can therefore be mimicked using such hydrogels to improve our understanding of how the fibrotic response is perpetuated by the feedback from the fibrotic microenvironment itself during IPF.

In this study, we hypothesized that an altered microenvironment in fibrotic lungs contributes to perpetuation of fibrosis by inducing profibrotic behavior of lung fibroblasts. To address this, we used an *in vitro* model using IPF and control lung ECM-derived hydrogels cultured with either IPF or control lung-derived primary fibroblasts in a combined fashion for 7 and 14 days. We investigated the influence of the fibrotic microenvironment on both IPF and control fibroblasts by comparing the ECM remodeling responses of the fibroblasts encapsulated in IPF and control ECM in 3D. We characterized fibroblast induced changes to the microenvironment with respect to modulation of collagen and glycosaminoglycan (GAG) content, collagen fiber organization and mechanical properties by comparing the fibroblast-driven ECM remodeling responses’ with empty hydrogels.

## Materials and Methods

### Experimental Design

The experimental approach adopted in this study is described in **Figure 1**. The specific details of the methods are described below.

**Figure 1:**
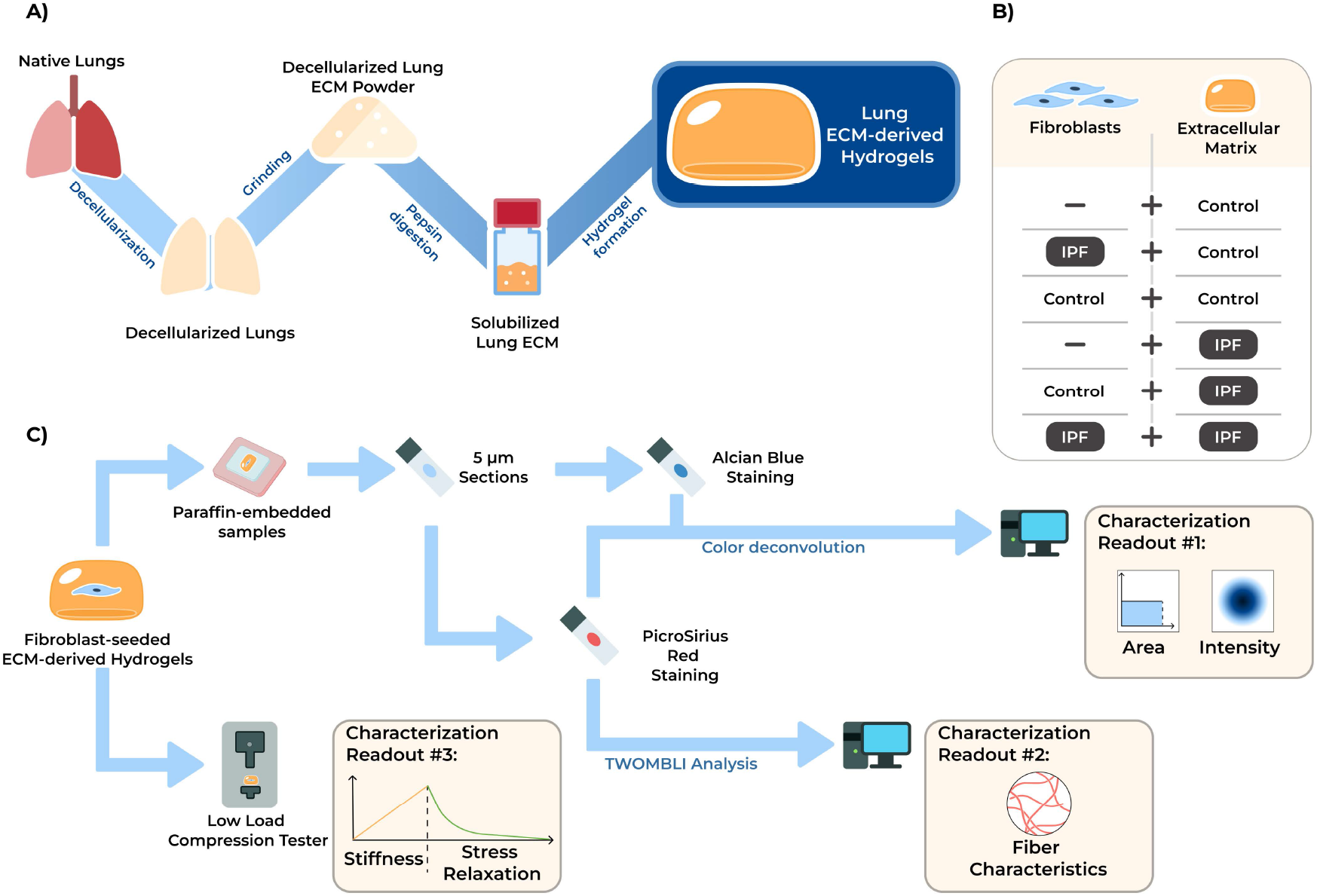
Overview of the experimental design and readouts measured in this study. A) Control and IPF lungs were decellularized, ground to a fine powder and digested using pepsin in acidic medium. The solubilized lung ECM was used to prepare lung ECM-derived hydrogels. B) Combinatorial approach used in the experiments. Fibroblasts and ECM groups were cross-combined to have combinations of control and IPF originated samples in every experimental batch. C) The readout applied in this study: PicroSirius Red and Alcian Blue stained sections of empty and fibroblast-encapsulated control and IPF lung ECM-derived hydrogels were scanned and digitally analyzed for area only or area and intensity (Characterization Readout #1). Fluorescence images of PicroSirius Red stained sections of empty and fibroblast-encapsulated control and IPF lung ECM-derived hydrogels were analyzed using TWOMBLI plugin in ImageJ to analyze the fiber characteristics (Characterization Readout #2). The mechanical properties (stiffness and stress relaxation) of empty and fibroblast-encapsulated control and IPF lung ECM-derived hydrogels were measured using low load compression tester (Characterization Readout #3).

### Lung decellularization

Decellularized control (macroscopically normal tissue, referred to as control throughout the manuscript) and lung tissue from patients with IPF were kindly provided by Dr. Steven Huang, University of Michigan, USA. De-identified control and IPF human lung tissue were provided by the University of Michigan; as the tissues were de-identified and coming from deceased donors, the University of Michigan Institutional Review Board deemed this work exempt from oversight. The decellularization procedure was performed as described previously [11]. Briefly, the fresh lung tissue samples were washed with 1X PBS (Gibco, Waltham, MA, United States). The tissue sample was washed for 24 hours per step at 4°C (unless otherwise stated) with the series of different solutions under constant agitation conditions: 1% (v/v) Triton X-100 (Sigma Aldrich, St. Louis, MO, USA), 2% (wt/v) sodium deoxycholate (Sigma Aldrich), 1 M NaCl (Sigma Aldrich), 30 mg/L DNase (Sigma Aldrich) with 1.3 mM MgSO_4_.7H_2_O (Sigma Aldrich) and 2 mM CaCl_2_ (Sigma Aldrich) at 37°C. The washing series was performed twice with three times PBS washes between every different solution. Afterwards, the ECM samples were treated with 0.18 % peracetic acid solution (32% w/w; Merck, Darmstadt, Germany) in 4.8% ethanol (Fresenius Kabi, Bad Homburg vor der Höhe, Germany) solution for 24 hours at 4°C with constant shaking. Lastly, the ECM samples were washed using PBS and kept in PBS (+1% Penicillin-Streptomycin (Gibco)) at 4°C until the next step.

### Lyophilization and grinding of decellularized lung samples

Decellularized lung samples were lyophilized in order to remove excess water. Briefly, the samples were snap-frozen using liquid nitrogen, then freeze-dried using a Labconco Freezone 2.5 Liter Benchtop Lyophilizer (Labconco, Kansas City, MO, USA) until the samples reached complete dryness. Once the lung scaffold samples were dry, they were brought to room temperature for the grinding process. An IKA A11 Basic Analytical Mill (IKA, Germany) was used to grind the lung scaffold pieces to a fine powder. Lung ECM powders were kept at room temperature with desiccant until use.

### Preparation of lung ECM-derived hydrogels

Decellularized lung ECM powders were pooled (n = 7 each for both control and IPF lung ECM samples) with equal amounts of powder dry weight per donor in order to minimize the patient-to-patient variation. Using this powder, pepsin digestion was performed using 2 mL of 2 mg/mL pepsin (Sigma-Aldrich) solution in 0.01 M HCl for 40 mg dry ECM powder in a 7.5 mL glass vial as previously described [25]. The digestion was performed for 72 hours with constant stirring at room temperature. After the incubation, the pH was brought back to physiological conditions (pH = 7.4) using 0.1 M NaOH (Sigma-Aldrich) solution before being mixed with 10X PBS (Gibco) solution to supplement the solution (will be referred to as pre-gel from here on) with physiological salts and ions. Resulting control and IPF pre-gels were cast into the wells of a 48-well plate or in a 1.5 mL Eppendorf tube and incubated for 1-2 hours to allow gelation and after the incubation, the hydrogel formation was checked.

### Primary lung fibroblast isolation and cell culture

Control and IPF primary human lung fibroblasts were isolated from lung tissue of patients undergoing surgery for tumor resection or lung transplantation at the University Medical Center Groningen (UMCG) as previously described [26]. Control tissue was obtained from macroscopically normal looking tissue from tumor excision surgery, as far as possible from the tumor. IPF primary human lung fibroblasts were isolated from the peripheral lung tissue of explanted lung tissue of patients who underwent lung transplantation as previously described. This study protocol was consistent with the Research Code of the University Medical Center Groningen (https://www.umcg.nl/documents/770534/2183586/umcg-research-code-2018-en.pdf/9680a460-3feb-543d-7d58-bc9d4f7277de?t=1614951313016, last accessed 18/07/2023) and the national ethical and professional guidelines (“Code of conduct for Health Research (only in Dutch):Gedragscode-Gezondheidsonderzoek-2022.pdf (https://www.coreon.org/gedragscode-gezondheidsonderzoek, last accessed 18/07/2023). Lung tissues used in this study were derived from leftover lung material after lung surgery from archival materials that are exempt from consent in compliance with applicable laws and regulations (Dutch laws: Medical Treatment Agreement Act (WGBO) art 458 / GDPR art 9/ UAVG art 24). This material was not subject to the Medical Research Human Subjects Act in the Netherlands, and, therefore, an ethics waiver was provided by the Medical Ethical Committee of the University Medical Center Groningen.

The characteristics of the patients are presented in **Table 1**. The fibroblast cultures were maintained in complete growth media (DMEM Low Glucose (Gibco), supplemented with 10% FBS (Sigma-Aldrich), 1% Penicillin-Streptomycin (Gibco) and 1% GlutaMAX (Gibco)) and only used when tested negative against *mycoplasma* infection using a PCR screening. Fibroblasts were used at passage 5.

**Table 1:** Patient characteristics of the fibroblasts donors used in this study. ^*^: tested with Mann-Whitney U test. ^†^: tested with Fisher’s exact test. ES: Ex-smoker, F: Female, IPF: Idiopathic pulmonary fibrosis, M: Male, NS: Non-smoker.

|  | Control | IPF | P value |
| --- | --- | --- | --- |
| Age (median, (min-max)) | 65, (59-72) | 63.5, (61-68) | 0.688 * |
| Sex (M/F) | 6/0 | 6/0 | >0.999<sup>†</sup> |
| Smoking status | 5 EX, 1 NS | 4 EX, 2 NS | >0.999<sup>†</sup> |

### Seeding primary lung fibroblasts in lung ECM-derived hydrogels

Primary human lung fibroblasts (n = 6 for both control and IPF, all at passage 5) were harvested using 0.25% Trypsin-EDTA (Gibco) and centrifuged at 500 x *g* for 5 minutes. Afterwards, the supernatant was discarded and the pellet resuspended using 10 mL full complete growth media to count the cells using an automated cell counter NC-200 (Chemometec, Denmark). Then, the cell suspension was transferred to new tubes at a concentration of 2.5 × 10^6^ cells per tube and centrifuged again. The supernatants were discarded and the pellets were resuspended using 2.5 mL pre-gel of each type. After ensuring the proper dispersal of the cell pellet in the pre-gel solution, 200 μL cell suspension was cast per well of 48-well plates and incubated for 2 hours. For each experimental set, empty hydrogels from control or IPF pre-gels were also cast and used as fibroblast-free controls. After observing hydrogel formation, the empty and fibroblast-encapsulated hydrogels were supplemented with 400 μL complete growth media for the culture period. The gels were incubated for 7 and 14 days with complete growth media in a CO_2_ incubator (5% CO_2_, 37°C), with half change of growth media at days 4, 7 and 11.

### Paraffin Embedding, Sectioning, and Deparaffinization

ECM-derived hydrogels (both fibroblast-encapsulated and empty) were fixed using 500 μL 2% paraformaldehyde (PFA; Sigma-Aldrich) for 30 minutes at room temperature after removing the growth media from the wells at the end of 7- or 14-day culture period. Afterwards, they were embedded in 1% agarose (Invitrogen, Waltham, MA, USA) solution to prevent dehydration. Agarose-embedded hydrogels then were fixed with 4% formalin and embedded in paraffin. Five μm sections from the paraffin-embedded samples were cut and placed on Star Frost (Knittel Glass, Braunschweig, Germany) glass slides and incubated at 65°C for 1 hour to ensure retention of the sections on the slides. Then, the slides were deparaffinized using serial ten minute incubations in xylene (Klinipath BV, Duiven, Netherlands) solution, 100% ethanol (Fresenius Kabi), 96% ethanol, 70% ethanol and distilled water.

### Picrosirius Red Staining for visualization of collagens

Collagens were visualized using PicroSirius Red (PSR), which binds to all collagen molecules through electrostatic interactions [27]. PSR solution was prepared using 0.5 g Sirius red F3B (Sigma) in 500 mL saturated aqueous picric acid solution. Deparaffinized slides were washed with distilled water and then incubated with the PSR solution for an hour. Afterwards, the slides were washed with acidified water (5 mL glacial acetic acid (Merck) in 11 mL distilled water) twice and dehydrated through 75%, 96%, 100% ethanol (Fresenius Kabi) and xylene (Klinipath BV) solutions. After airdrying the slides, they were mounted using a non-aqueous mounting medium and kept in dark until imaging.

### Alcian Blue Staining for visualization of acidic glycosaminoglycans

Alcian blue solution was prepared using 1 g Alcian blue (Sigma-Aldrich) powder in 100 mL 3% acetic acid solution. Nuclear fast red solution was prepared using 0.1 g nuclear fast red powder in 100 mL distilled water with 5 g Al_2_(SO4)_3_ (Sigma-Aldrich). Deparaffinized slides were washed with distilled water and then incubated with the Alcian blue solution for 30 minutes at room temperature. Afterwards, the slides were washed with running tap water for 2 minutes and rinsed in distilled water. Counter-staining was performed through incubation with nuclear fast red solution for 5 minutes and the slides were washed in running tap water for 1 minute. The slides then were dehydrated through 75%, 96%, 100% ethanol (Fresenius Kabi) and xylene (Klinipath BV) solutions. After airdrying the slides, they were mounted using a non-aqueous mounting medium and kept in dark until imaging.

### Imaging

Immunohistochemical staining results were imaged using a Hamamatsu scanner (Hamamatsu Photonics K.K., Herrsching, Germany) at 40X magnification. Fluorescent microscopy images of the PSR stained hydrogels were captured with Leica SP8X white light laser confocal microscope (Leica, Wetzlar, Germany) with UV-vis absorption (λ_ex_) 561 nm and emission (λ_em_) 566 / 670 nm using 63x/1.40 Oil immersion lens with a digital zoom 2X.

## Image Analysis

### Analysis of Light Microscopy Images

Specific image areas with stained gel were extracted into TIF (LZW) files using Aperio ImageScope V12.4.0.5043 (Leica Biosystems, Amsterdam, Netherlands). These TIF files were opened in Adobe Photoshop CS6 Extended (San Jose, California, USA) and artifacts, such as obvious background staining, hairs or other contaminations and folded areas of gels, were removed. Ten areas from ten distinct regions per image (1 image per gel) were randomly selected and saved as separate image files to enable calculation of the average lowest threshold for identifiable specific staining for all images. These 10 images were opened in FIJI 1.53F51 (LOCI, University of Wisconsin) [28] as 8-bit images and split into different channels that captured the different staining colors, using the appropriate vector for the thresholds of the staining using color deconvolution. An in-house built macro (*Supplementary Document 1* for PSR Images and *Supplementary Document 2* for AB Images) was used along with the FIJI SlideJ plugin (tile size: 20000) to calculate the outputs from all images. Results were saved as a text to tab file and opened in R studio 4.1.1 (Boston, MA, USA) to sort the percentage area of positively stained pixels and the average intensity of the positively stained pixels data of the different staining levels (Supplementary Figure 1) using an in-house built macro as previously described [29]. Briefly, the strength of the signal from each pixel was determined and categorized as “weak”, “moderate” or “strong” based on the level of the strength. The pixels that belong to each of these categories were then combined per image and used in the following steps. Area percentage and intensity values for each category per image were calculated in Microsoft Excel 2016 (Microsoft, Resmond, WA, USA). For these calculations, the sorted rows Area_Colour (area) and RawIntDen_Colour (intensity) were used. Stained area (%) and intensity values (arbitrary units) of each category was calculated using equation (i) and equation (ii), respectively, as shown below.

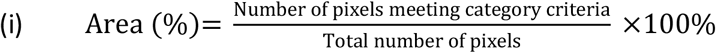

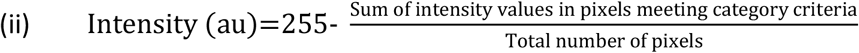

### Analysis of Fluorescence Microscopy Images

For every sample, images were generated of 6 randomly selected areas and saved as TIF (LZW) files. Each image was analyzed using the The Workflow Of Matrix BioLogy Informatics (TWOMBLI) plugin in FIJI 1.53F51 (LOCI) [30].: For the global fiber parameters, the following parameters were examined: the area, percentage of high density matrix, which shows percentage area covered by fibers that are detected to have a highly dense arrangement (many fibers within a small area) based on pixel saturation, and alignment of fibers, which denotes the percentage of fibers with similar orientation. For the individual fiber parameters, the following parameters were examined: total fiber length, end points, branchpoints. The curvature analysis was performed using curvature windows 20 and 50 in TWOMBLI plugin to investigate low and high curvature windows, respectively, to capture the individual wavinessof the fibers (how widely spread are the points at which the components of the curve cross a center line) through low curvature window analysis and changes in the peak height of the curves through high curvature window analysis.

### Mechanical testing with Low Load Compression Testing (LLCT)

Stiffness values and viscoelastic relaxation properties of the hydrogels at day 7 and 14 were measured using a Low Load Compression Tester as previously described [31-33]. The LLCT analysis was performed on three different randomly selected locations on each hydrogel using 20% fixed strain rate. The measurement locations had at least 2 mm distance between them and 2 mm from the edges to ensure robustness and representativeness of the measurements. The stress ((Equation (iii)) and strain (Equation (iv)) values were calculated from the linear elastic region as described below and the slope of the line was used to calculate Young’s modulus (E, stiffness) (Equation (v)) until the peak point for the highest measurement observed). Relaxation values were calculated starting from the time point at which highest stiffness was observed by using the formula (Equation (vi)). Representative stress-strain curves and stress relaxation profiles for each group are illustrated in *Supplementary Figures 2&3*. Time duration for reaching 100% relaxation was recorded as ‘Relaxation time’. Time duration for reaching 50% of the total relaxation was recorded as ‘Time to Reach 50% Stress Relaxation’. All calculations were performed using Microsoft Excel 2016.

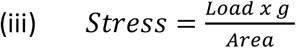

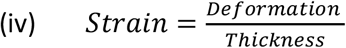

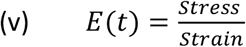

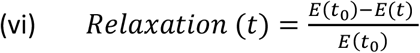

## Statistical Analysis

Statistical analyses of the measured parameters were performed using an interaction analysis in a mixed model analysis in IBM SPSS Statistics 26 (IBM, Armonk, New York, USA). For each parameter analyzed, the interactions between the disease status of the hydrogels and disease status of the fibroblasts, as well as fibroblast encapsulation status of the hydrogels were used. **Figure 2** illustrates the analysis strategy performed. For all analyses presented, a random effect was used for the intercept per experimental batch (i.e. same combination of control or IPF fibroblasts and control or IPF hydrogel for the 6 experimental conditions (n=6 batches)). TWOMBLI results were analyzed using 6 different images generated per sample to address the sample heterogeneity. Mechanical characterization results were analyzed using triplicated measurements performed on the same sample to tackle the heterogeneity within samples. Presented results show estimate ± 95% confidence interval for all results.

**Figure 2:**
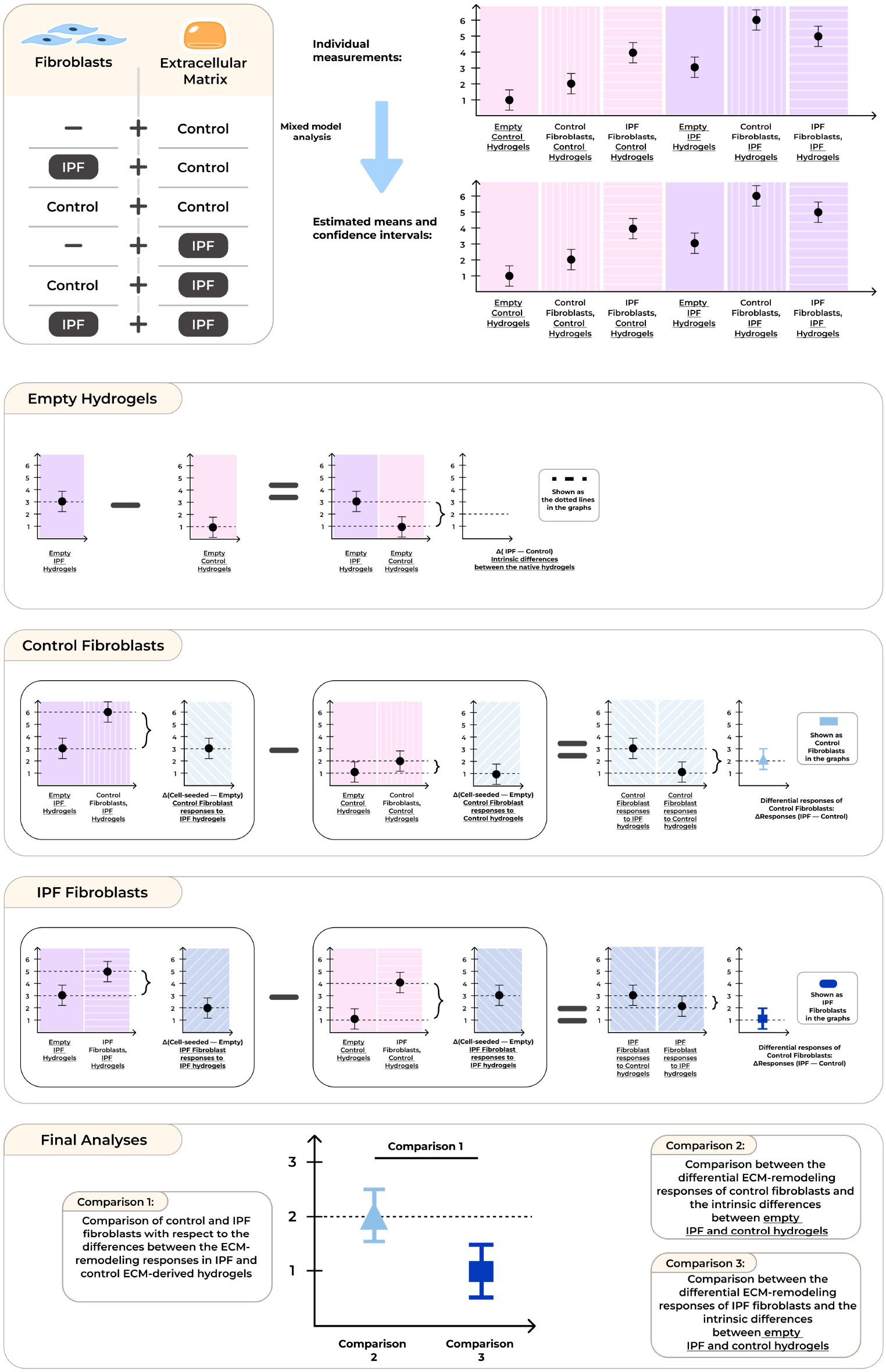
Analysis approach used throughout the study. For each measurement, six IPF and control lung donor-derived fibroblasts were encapsulated in both types of hydrogels were used as biological replicates and compared with empty hydrogels prepared in the same experimental batch. A mixed-model analysis was used to evaluate the statistical differences between the groups. The differences between the empty and fibroblast-encapsulated hydrogels were calculated for both IPF and control ECM-derived hydrogels. The differential ECM-remodeling responses of the fibroblasts with respect to the measurement performed were calculated by subtracting the fibroblast-originated ECM-remodeling responses measured in control hydrogels from those measured in IPF hydrogels.

## Results

### Fibroblasts change collagen organization only in IPF hydrogels

We first assessed the impact of interactions between a fibrotic microenvironment and fibroblasts on totalcollagen presence in the lung ECM-derived hydrogels by analyzing the collagen content at day 7 and 14. PicroSirius Red (PSR)-stained sections of empty or fibroblast-encapsulated control and IPF hydrogels were analyzed using an automated image analysis that separated and then compared the amount of collagen present in the hydrogesl at three different levels of pixel strength: weak, moderate and strong (*Supplementary Figure 1*). In all three categoriesof pixel strengths, we did not detect any differences in the percentage of PSR-stained area present among the different groups (*Supplementary Figure 4, Supplementary Tables 1-6*). Similarly, the mean intensities of staining compared within the weak, moderate or strong pixel strengths of PSR-staining were not different between the tested groups (*Supplementary Figure 5, Supplementary Tables 7-12*). We also examined the distribution of GAGs in all groups by staining sections with Alcian Blue. Comparable to collagen content, the pecentage area of GAG content was similar in control and IPF empty hydrogels, and this did not change in hydrogels in which control or IPF fibroblasts were encapsulated (*Supplementary Figure 3, Supplementary Tables 13-18*).

We then investigated whether fibroblasts induced changes in collagen organization and whether this was different in fibrotic versus control hydrogels using TWOMBLI. This analysis revealed the changes triggered in fibrotic ECM (**Figure 3**). The first parameter we studied was the percentage area occupied by high density matrix, which describes the collagen fiber organization at a global level (**Figure 3A**). Intrinsic differences between control and IPF hydrogels were not detected on day 7, but were clearly visible on day 14 with IPF hydrogels having more high density matrix than control hydrogels (*Supplementary Tables 19&20*, also denoted as the dotted lines in **Figure 3B-C)**. When we encapsulated fibroblasts in these hydrogels, we found that control fibroblasts further decreased the percentage of high density matrix in IPF hydrogels compared to control hydrogels below the already existing differences, both on day 7 and day 14 (**Figure 3B and 3C**, respectively). On the other hand, IPF fibroblasts decreased the percentage of high density matrix of IPF hydrogels only on day 14 (**Figure 3C**). The differential modulation of the hydrogels by control and IPF fibroblasts was significantly different (p = 5.12 × 10^−6^ for day 7 and p = 0.027 for day 14), indicating that not only hydrogel type but also the fibroblast origin contributes to dysregulated collagen organization in IPF.

**Figure 3:**
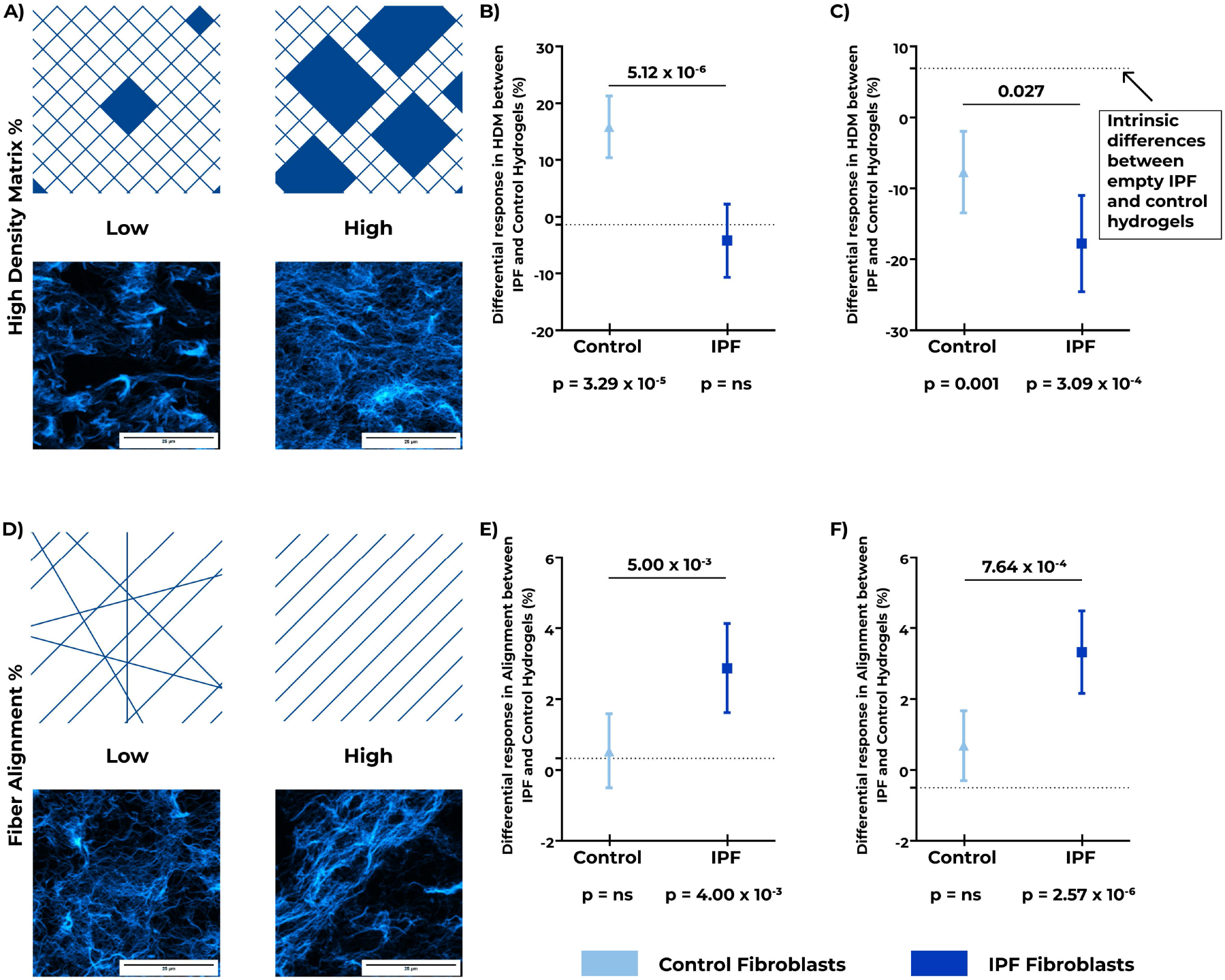
Changes in the global fiber organization in empty and fibroblast-encapsulated control and IPF lung ECM-derived hydrogels. Control and IPF primary lung fibroblasts were encapsulated in control or IPF lung ECM-derived hydrogels and cultured for 7 or 14 days. Fluorescence images of PicroSirius Red (PSR) stained sections of paraffin-embedded hydrogels were analyzed for global fiber characteristics and compared with their corresponding empty hydrogel samples. A) Schematic representation and example fluorescence images of high and low percentages of high-density matrix (HDM). B) Day 7 fibroblast-driven ECM remodeling response analysis for HDM (% area), C) Day 14 fibroblast-driven ECM remodeling response analysis for HDM (% area), D) Schematic representation and example fluorescence images of high and low percentages of fiber alignment. E) Day 7 fibroblast-driven ECM remodeling response analysis for fiber alignment (% fibers), F) Day 14 fibroblast-driven ECM remodeling response analysis for fiber alignment (% fibers). The dotted line shows the intrinsic difference between the <u>empty</u> IPF and control hydrogels. The estimate (± 95% confidence interval) shows the difference between the IPF and control hydrogels laden with control (light blue, triangle) or IPF (dark blue, square) fibroblasts. P values below each fibroblast group represent the differences induced by fibroblasts in IPF versus control hydrogels compared to the intrinsic difference between IPF and control empty hydrogels. P-values above the estimates indicate the differences between the responses of IPF and control fibroblasts in the different hydrogels. Applied statistical test: mixed-model analysis. ns: not significant, IPF: Idiopathic pulmonary fibrosis. n=6 for fibroblast donors, 6 images per sample were captured and analyzed.

We then characterized the degree of fiber alignment within the hydrogels (**Figure 3D**). We did not detect any intrinsic differences between control and IPF hydrogels with respect to fiber alignment (*Supplementary Tables 21&22*). When control fibroblasts were encapsulated in each type of hydrogel, they did not change the percentage fiber alignment in IPF hydrogels compared to control hydrogels at any time point (**Figure 3E** and **3F**). However, when encapsulating IPF fibroblasts, we found fiber alignment was greater in IPF hydrogels compared to control hydrogels on both day 7 and day 14 (**Figure 3E** and **3F**, p = 4.00 × 10^−3^ for day 7 and p = 2.57 × 10^−6^ for day 14). The differential modulation of the fiber alignment by control and IPF fibroblasts was significantly different on both days (p = 5.00 × 10^−3^ for day 7, p = 7.64 × 10^−4^ for day 14).

### Fibroblasts modify individual collagen fibers differently in fibrotic compared to control ECM

In addition to global fiber organization, individual fiber structure could also be altered by the encapsulated fibroblasts. Individual fiber structure was therefore assessed using TWOMBLI. Three individual fiber structural parameters were analyzed: average fiber length (AFL, μm), number of endpoints per 1000 μm fiber total length, and number of branchpoints per 1000 μm fiber total length (**Figure 4**).

**Figure 4:**
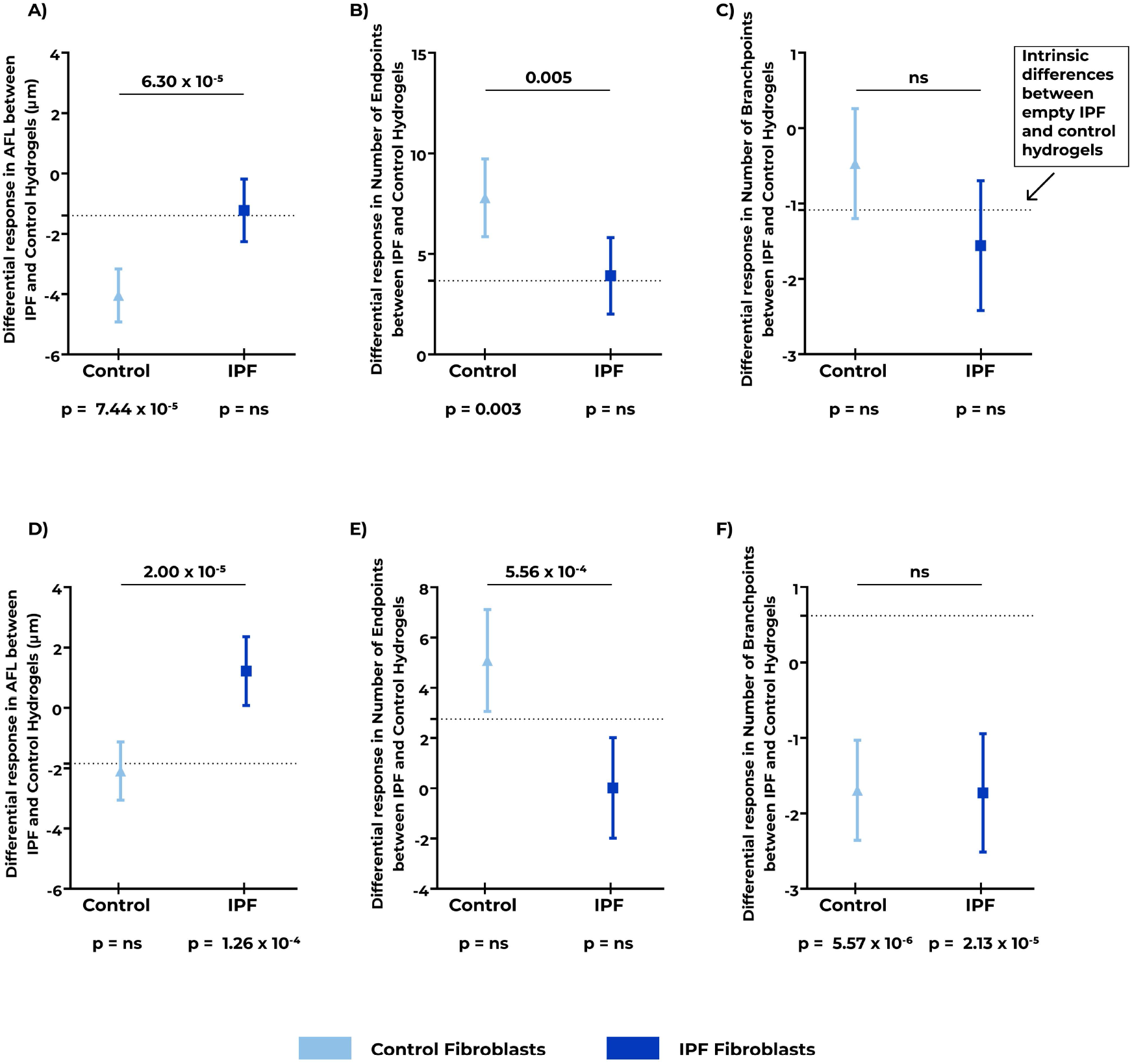
Individual collagen fiber structure in empty and fibroblast-encapsulated control and IPF lung ECM-derived hydrogels. Control and IPF primary lung fibroblasts were encapsulated in control or IPF lung ECM-derived hydrogels and cultured for 7 or 14 days. Fluorescence images of PicroSirius Red (PSR) stained sections of paraffin-embedded hydrogels were analyzed for individual fiber characteristics and compared with their corresponding empty hydrogel samples. <u>Day 7 fibroblast-driven ECM remodeling responses</u> with respect to A) average fiber length (AFL) (μm), B) number of endpoints per 1000 μm fiber total length, C) number of branchpoints per 1000 μm fiber total length. <u>Day 14 fibroblast-driven ECM remodeling responses</u> with respect to D) average fiber length (AFL) (μm), E) number of endpoints per 1000 μm fiber total length, F) number of branchpoints per 1000 μm fiber total length. The dotted line shows the intrinsic difference between <u>empty</u> IPF and control hydrogels. The estimate (± 95% confidence interval) shows the difference between the IPF and control hydrogels laden with control (light blue, triangle) or IPF (dark blue, square) fibroblasts. P values below each fibroblast group represent the differences induced by fibroblasts in IPF versus control hydrogels compared to the intrinsic difference between IPF and control empty hydrogels. P-values above the estimates indicate the differences between the fibroblast-driven ECM remodeling responses of IPF and control fibroblasts in the different hydrogels. Applied statistical test: mixed-model analysis. ns: not significant. IPF: Idiopathic pulmonary fibrosis. n=6 for fibroblast donors, 6 images per sample were captured and analyzed.

The analysis of average fiber length illustrated that, intrinsically, IPF hydrogels had shorter fibers on average, compared to control hydrogels (*Supplementary Tables 23&24*, also shown as the dotted line in **Figure 4A** and **4D** for day 7 and 14, respectively). When control fibroblasts were encapsulated in IPF hydrogels, the average fiber length was further decreased in IPF hydrogels on day 7 (**Figure 4A**, p = 7.44 × 10^−5^), while this modulation was not detected on day 14 (**Figure 4D**). IPF fibroblasts, on the other hand, increased the average fiber length in IPF hydrogels compared to control hydrogels only on day 14 (**Figure 4D**, p = 1.26 × 10^−4^). These modulations by control and IPF fibroblasts were significantly different from each other both on day 7 (p = 6.30 × 10^−5^) and day 14 (p = 2.00 × 10^−5^).

The number of individual fibers (number of endpoints) was higher in IPF hydrogels compared with control hydrogels on day 7, but this difference was not detected on day 14 (*Supplementary Tables 25&26*, also shown as the dotted line in **Figure 4B** and **4E** for day 7 and 14, respectively). When control fibroblasts were encapsulated in these hydrogels, we found more endpoints in IPF hydrogels compared to control hydrogels on day 7, above the existing intrinsic difference between these two hydrogels (**Figure 4B**). This modulation, however, was not observed on day 14 (**Figure 4E**). In contrast, IPF fibroblasts did not change the number of endpoints in IPF hydrogels compared to control hydrogels on either day 7 or day 14. These differential fibroblast-driven ECM remodeling responses between control and IPF fibroblasts, differed significantly on both day 7 (p = 0.005) and day 14 (p = 5.56 × 10^−4^) (**Figure 4B** and **4E**, respectively).

We also investigated the number of branchpoints as a measure of the number of fibers that had connections with other fibers. IPF hydrogels had more branchpoints than control hydrogels on day 7 (p = 0.007) while this intrinsic difference was not detectable on day 14 (*Supplementary Tables 27&28*, also shown as the dotted line in **Figure 4C** and **4F** for day 7 and 14, respectively). When either control or IPF fibroblasts were encapsulated in these hydrogels, no additional changes in the number of branchpoints induced by fibroblasts were detected in either control or IPF hydrogels on day 7 (**Figure 4C**). However, on day 14 both control (p = 5.57 × 10^−6^) and IPF fibroblasts (p = 2.13 × 10^−5^) strongly decreased the number of branchpoints in IPF hydrogels but not in control hydrogels (**Figure 4F**).

### Fibrotic microenvironment triggered differential regulation of fiber curvature by fibroblasts

We then moved on to characterize changes in the curvature of the collagen fibers in empty and fibroblast-encapsulated ECM-derived hydrogels as another parameter for comparing ECM remodeling responses of control and IPF fibroblasts to their microenvironment. The curvature of fibers was analyzed with respect to low and high curvature windows (**Figure 5A**). Low curvature windows capture the individual waviness (periodicity) of the fibers (more micro-scale) while higher curvature windows detect the global changes in the fiber shapes (changes in the peak height of the curves). These parameters are useful for describing the topographical arrangement of the fibers within the ECM hydrogel. Intrinsic differences between IPF and control hydrogels in the low curvature window were present in day 7 samples (p = 0.004), while they were not apparent for the high curvature windows (Supplementary Table 29&30, respectively, also shown as the dotted lines in **Figure 5B**). Control fibroblasts encapsulated in IPF hydrogels increased the fiber periodicity in low curvature windows (p = 0.001, **Figure 5B**) while they did not change the curvature heights in the higher curvature windows. IPF fibroblasts did not induce any periodicity changes (as measured in the low curvature windows); however, these fibroblasts reduced the peak heights of the fiber curves (as measured in the high curvature windows) in IPF hydrogels beyond the existing differences between IPF and empty hydrogels (p = 0.049, **Figure 5B**). Both low and high curvature window ECM remodeling responses of control and IPF fibroblasts to fibrotic hydrogels differed from each other (p = 3.2 × 10^−4^ for low and p = 0.006 for high curvature windows). Empty hydrogels analyzed on day 14 revealed intrinsic differences between IPF and control hydrogels in low (p = 1.15 × 10^−11^), and high (p = 0.026) curvature window samples (Supplementary Table 31&32, respectively, also shown as dotted line in **Figure 5C**). The existing differences between the IPF and control hydrogels were decreased by both control (p = 3.57 × 10^−4^) and IPF (3.01 × 10^−10^) fibroblasts in low curvature window. While both groups of fibroblasts decreased the fiber curvature periodicity in IPF hydrogels compared to control hydrogels, their ECM remodeling responses significantly differed from each other as well (p = 7.74 × 10^−4^). On the other hand, only IPF fibroblasts responded to fibrotic hydrogels in measurements in the high curvature window (p = 0.002). Even though only IPF fibroblasts altered the fiber curvature height, there were no apparent differences in the responses of IPF and control fibroblasts in this setting. These data indicate that IPF fibroblasts altered the topographical arrangement of the collagen fibers within their 3D microenvironment to a greater extent than the control fibroblasts.

**Figure 5:**
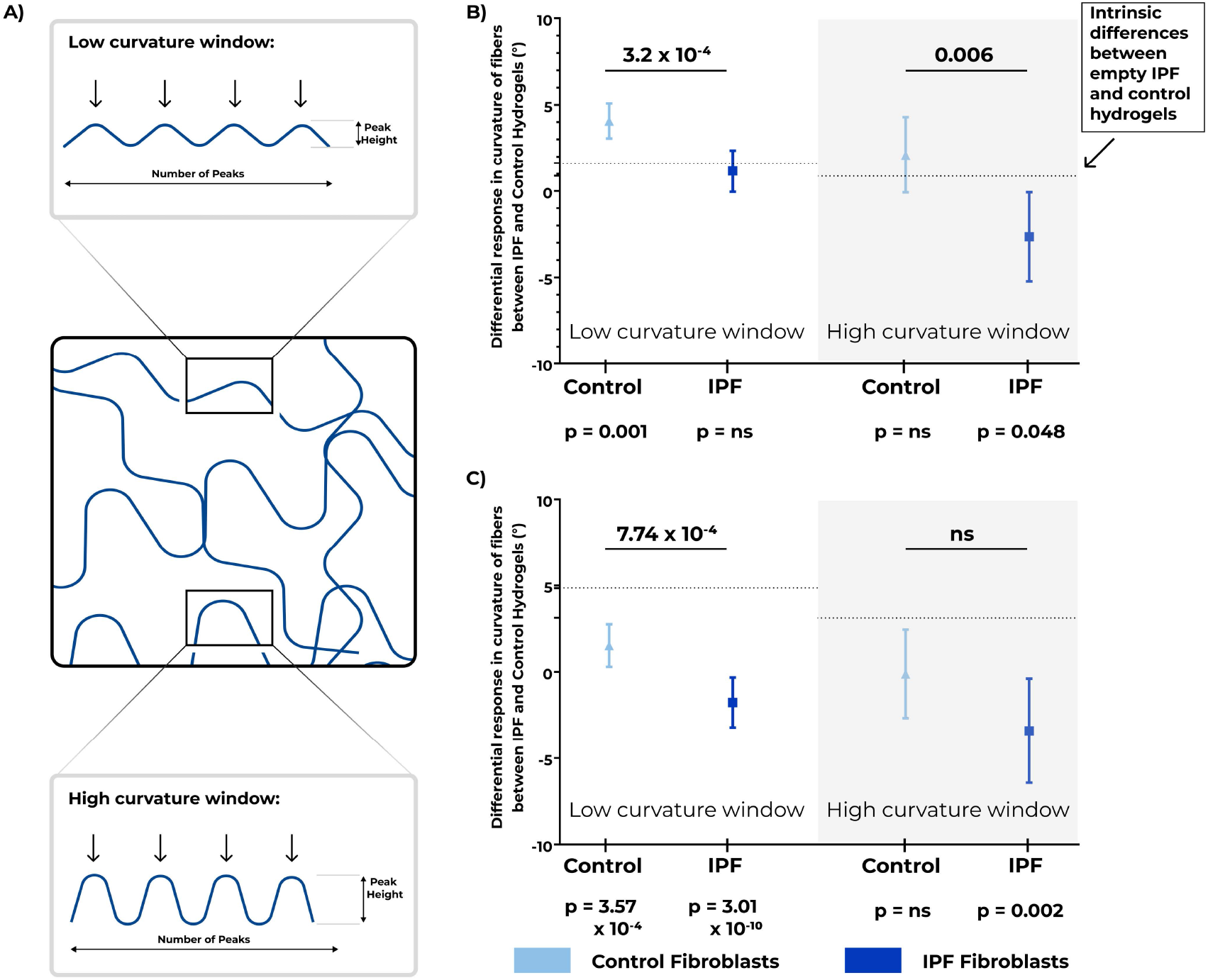
Investigation of collagen fiber curvature in empty and fibroblast-encapsulated control and IPF lung ECM-derived hydrogels. Control and IPF primary lung fibroblasts were encapsulated in control or IPF lung ECM-derived hydrogels and cultured for 7 or 14 days. Fluorescence images of PicroSirius Red (PSR) stained sections of paraffin-embedded hydrogels were analyzed for the differences in collagen fiber curvature and compared with their corresponding empty hydrogel samples. A) Schematic representation of low and high curvature windows with respect to periodicity of peaks and peak height, B) <u>Day 7</u> fibroblast-driven ECM remodeling response analysis for fiber curvature with respect to number of peaks and peak height, C) <u>Day 14</u> fibroblast-driven ECM remodeling response analysis for fiber curvature with respect to number of peaks and peak height. The dotted line shows the intrinsic difference between <u>empty</u> IPF and control hydrogels. The estimate (± 95% confidence interval) shows the difference between the IPF and control hydrogels laden with control (light blue, triangle) or IPF (dark blue, square) fibroblasts. P values below each fibroblast group represent the differences induced by fibroblasts in IPF versus control hydrogels compared to the intrinsic difference between IPF and control empty hydrogels. P-values above the estimates indicate the differences between the fibroblast-driven ECM remodeling responses of IPF and control fibroblasts in the different hydrogels. Applied statistical test: mixed-model analysis. ns: not significant, IPF: Idiopathic pulmonary fibrosis. n=6 for fibroblast donors, 6 images per sample were captured and analyzed.

### Stiffness increased but stress relaxation remained comparable when fibroblasts were in fibrotic hydrogels

Lastly, we characterized mechanical properties of both empty and fibroblast-encapsulated control and IPF hydrogels after 7 or 14 days of culture. Stiffness (**Figure 6A and 6B**) and viscoelastic stress relaxation (**Figure 6E and 6F**) behavior of hydrogels were measured using a Low-Load Compression Tester (LLCT). Empty IPF hydrogels were stiffer than empty control hydrogels on both day 7 (p = 5.57 × 10^−6^) and day 14 (p = 4.93 × 10^−22^) (*Supplementary Tables 33&34*, also shown as the dotted line in **Figure 6C** and **6D** for day 7 and 14, respectively), reflecting previous reports [12]. When control fibroblasts were encapsulated in IPF hydrogels, they stiffened the hydrogels significantly more than they stiffened control hydrogels on day 7 (**Figure 6C**, p = 0.002). Similar to control fibroblasts, IPF fibroblasts also increased stiffness of IPF hydrogels more than they stiffened control hydrogels (**Figure 6D**, p = 0.002). The ECM remodeling responses of control and IPF fibroblasts in IPF hydrogels compared to control hydrogels were analogous. However, on day 14 only control fibroblasts increased the stiffness of IPF hydrogels compared to control hydrogels beyond the already existing stiffness differences (**Figure 6D**, p = 9.22 × 10^−5^). Modulation of IPF hydrogels compared to control hydrogels by control fibroblasts on day 14 was significantly different from how IPF fibroblasts modulated the two types of hydrogels (p = 0.027).

**Figure 6:**
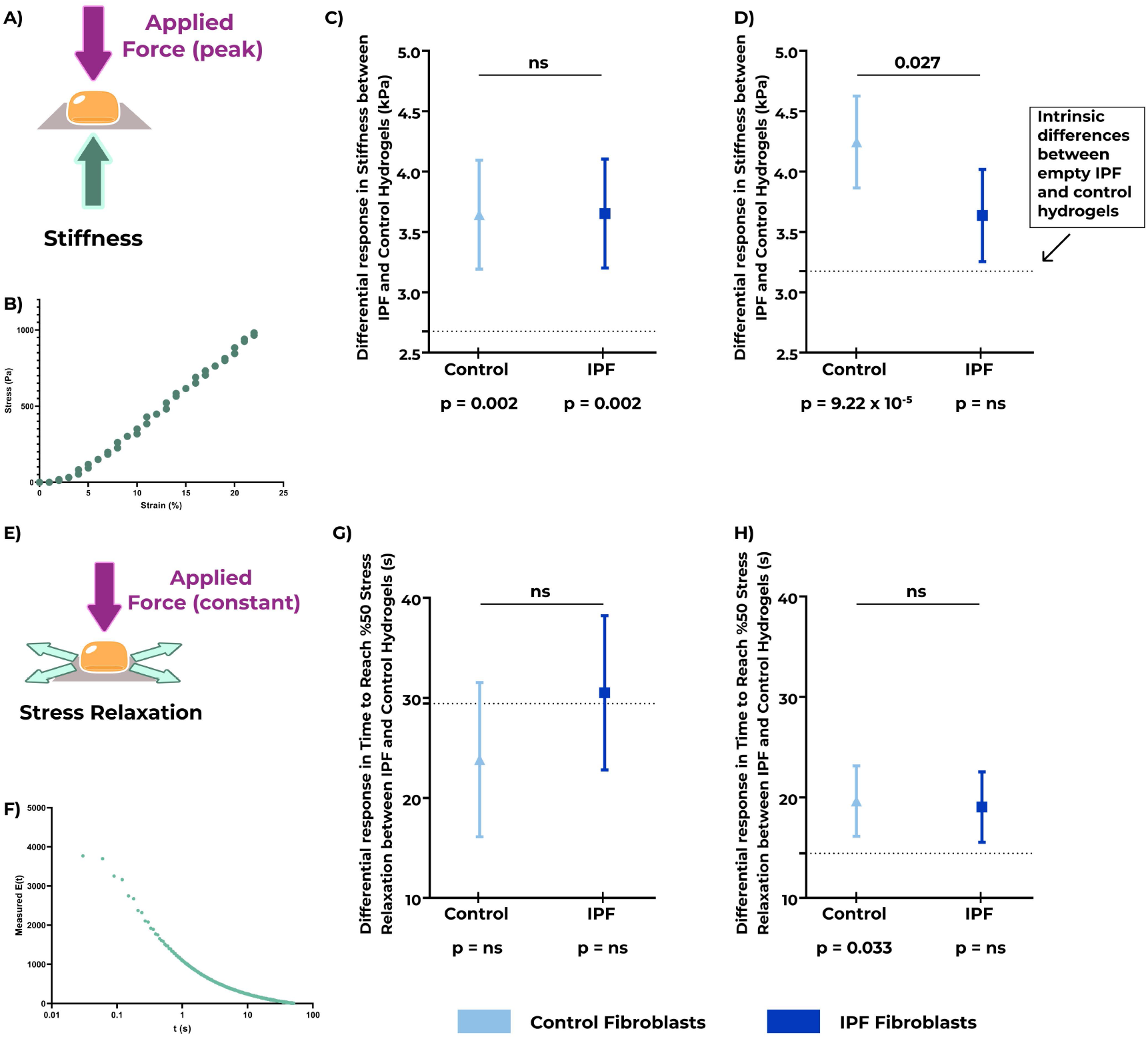
Mechanical properties in empty and fibroblast-encapsulated control and IPF lung ECM-derived hydrogels. Control and IPF primary lung fibroblasts were encapsulated in control or IPF lung ECM-derived hydrogels and cultured for 7 or 14 days. Fibroblast-encapsulated hydrogels were mechanically tested using low load compression testing (LLCT) and compared with their corresponding empty hydrogels. A) Schematic representation of the stiffness analysis performed using compression. B) An example stress-strain curve obtained using LLCT, C) <u>Day 7</u> fibroblast-driven ECM remodeling response analysis for stiffness of the hydrogels (kPa), D) <u>Day 14</u> fibroblast-driven ECM remodeling response analysis for stiffness of the hydrogels (kPa), E) Schematic representation of the stress relaxation analysis performed using compression. F) An example relaxation profile obtained using LLCT, G) <u>Day 7</u> fibroblast-driven ECM remodeling response analysis for time to reach 50% stress relaxation (s), H) <u>Day 14</u> fibroblast-driven ECM remodeling response analysis for time to reach 50% stress relaxation (s). The dotted line shows the intrinsic difference between <u>empty</u> IPF and control hydrogels. The estimate (± 95% confidence interval) shows the difference between the IPF and control hydrogels laden with control (light blue, triangle) or IPF (dark blue, square) fibroblasts. P values below each fibroblast group represent the differences induced by fibroblasts in IPF versus control hydrogels compared to the intrinsic difference between IPF and control empty hydrogels. P-values above the estimates indicate the differences between the fibroblast-driven ECM remodeling responses of IPF and control fibroblasts in the different hydrogels. Applied statistical test: mixed-model analysis. All measurements were performed at 20% strain rate. Applied statistical test: mixed-model analysis. ns: not significant, IPF: Idiopathic pulmonary fibrosis. n=6 for fibroblast donors, 3 independent locations per sample were measured.

Viscoelastic stress relaxation was the other mechanical parameter analyzed after 7 and 14 days of culture of control and IPF fibroblasts in control and IPF hydrogels. The time to reach 50% stress relaxation was measured in seconds and compared between the groups (**Figure 6E**). IPF hydrogels relaxed significantly slower than control hydrogels both on day 7 (p = 1.47 × 10^−12^) and day 14 (p = 5.82 × 10^−14^) (*Supplementary Tables 35&36*, also shown as the dotted line in **Figure 6G** and **6H** for day 7 and 14, respectively). Seeding of either control or IPF fibroblasts into the hydrogels did not significantly change stress relaxation beyond the intrinsic differences that were already present on day 7 (**Figure 6G**). On day 14, however, only control fibroblasts caused a slower relaxation of IPF hydrogels compared to control hydrogels, in addition to the existing differences (**Figure 6H**, p = 0.033). IPF fibroblasts did not change the relaxation properties of the hydrogels beyond the existing differences between control and IPF.

## Discussion

In this study, we used a 3D *in vitro* model composed of human lung ECM and human primary lung fibroblasts to assess the influence of the microenvironment on fibroblast-driven ECM remodeling responses. We showed that while collagen content and GAG content were unchanged by the fibroblast groups, fiber organization directed by the fibroblasts differed substantially due to influence of the ECM microenvironment. When examining individual collagen fiber characteristics the fibrotic microenvironment did not induce significant changes. However, the global structure of the ECM arrangement, as illustrated by fiber alignment and high density matrix proportion was impacted by the nature of the microenvironment. Control fibroblasts did not alter the fiber alignment, while fibrotic fibroblasts increased fiber alignment and this happened to a greater extent in the fibrotic microenvironment. In contrast, control fibroblasts modulated the percentage of high density matrix in a temporal manner in the fibrotic microenvironment, while the IPF fibroblasts reduced the percentage of high density matrix. In addition, control fibroblasts altered the topographical arrangement of the collagen fibers, giving them a greater degree of curvature than that seen with the IPF fibroblasts. The mechanical characteristics of the fibrotic microenvironment were increased by fibroblasts from both control and IPF donors, whereas this change was not seen in the control hydrogels. These findings illustrate that the fibrotic microenvironment imparts a powerful message that drives cellular responses. Overall, our results illustrate that the fibroblast-encapsulated lung ECM-derived hydrogel model is a powerful *in vitro* tool for understanding cell interactions with the local microenvironment, and divulging greater knowledge of feedback by the fibrotic ECM and how fibroblasts remodel the microenvironment during this response.

Collagen amount and organization are known to be drastically altered in IPF; higher amounts of collagen with an increased disorganization of the fiber structure have been consistently documented [8, 11, 13]. In our hydrogel system, fibroblasts did not induce changes in collagen amount between empty and fibroblast-encapsulated hydrogels, suggesting that these fibroblasts did not deposit detectable new collagen in this model and timeframe. A recent study reported an increase in protein levels of collagen types VII, X and XIV, detected using mass spectrometry, by control fibroblasts cultured in spheroid form with presence of IPF lung ECM, compared to non-IPF ECM [34]. These data provide further evidence, supporting prior reports [11, 14, 35, 36] that the IPF ECM provides a pro-fibrotic signal for fibroblasts. While our data appear to contrast the previous studies, the use of mass spectrometry may provide additional sensitivity that would enable a more penetrating investigation of the collagen changes. In our model system, the fibroblast-induced differences in high density matrix and collagen fiber alignment as directed by the type of environment (hydrogel) in which the cells were grown, implies that the lack of detectable changes in global total collagen amount does not necessarily reflect a lack in pro-fibrotic responses by fibroblasts.

Increased fiber density was previously proposed as a mechanism for triggering activation of fibroblasts [15]. Therefore, in a fibrotic microenvironment, with higher amounts of dense fibers [37, 38], greater fibroblast responses would be expected. Interestingly, control fibroblasts appeared to have opposite ECM remodeling responses at different time points, with an initial increase in high density matrix after 7 days but a subsequent decrease after 14 days. These differences in time might reflect how naïve fibroblasts are imprinted by a fibrotic microenvironment as time passes so their ECM remodeling responses induced by the fibrotic environment are changed. In IPF fibroblasts, the decrease in high density matrix with a concomitant increase in fiber alignment, points at exaggerated ECM remodeling responses of these fibroblasts, when confronted with fibrotic ECM. These specific responses of IPF fibroblasts suggest that the origin of both microenvironment and fibroblasts play a crucial role in determining the organizational changes of collagen fibers.

Enhanced collagen crosslinking is recognized to be enhanced in IPF lung tissues [9, 39]. Our results showing decreased amounts of high density matrix in fibrotic ECM modulated by both types of fibroblasts in day 14 samples do not initially seem to be in concert with previous reports showing enhanced fibroblast activation by increased ECM crosslinking in IPF [39]. Further research is required to examine if the high density matrix in this hydrogel model is composed of crosslinked collagen fibers or if it is non-covalent aggregations of fibers. The latter may have different cellular signaling implications than the highly cross-linked ECM in IPF tissue, parallel to the previous reports showing different levels of myofibroblast activation in chemically crosslinked hydrogels compared with physically crosslinked hydrogels [40]. While it was out of the scope of this study, it is important to recognize that a role of proteoglycans in collagen crosslinking and fiber organization has been previously reported [41], and their presence might also play a role in directing the collagen organization measured in this model.

The organization of individual collagen fibers is also an important element for determining cellular responses to the microenvironment in which they reside [11]. With respect to the individual fiber organization parameters that were analyzed in our study, control and IPF fibroblasts responded differently from each other, with the exception of the alterations to the number of branchpoints. The opposite ECM remodeling responses elicited by the fibroblasts to fibrotic and control hydrogels highlights that not only the origin of the microenvironment but also the origin of fibroblasts plays a role in dictating the collagen organization. Fiber curvature (collagen topographical arrangement), as both an individual and global fiber parameter, was differentially regulated by control and IPF fibroblasts encapsulated in fibrotic hydrogels compared to control hydrogels. It is intriguing that these changes coincided with the changes in the mechanical parameters initiated by the fibroblasts. In particular, control fibroblasts had an exaggerated response to the fibrotic microenvironment, resulting in an increased stiffness compared to control hydrogels. While the decrease in high density matrix and increase in stiffness do not seem to go hand-in-hand, fibroblasts might be realigning the fibers that are dissociated from the high density matrix in a manner that leads to the increased stiffness. The lack of changes in the peak height of the collagen fiber curves in control fibroblast-encapsulated hydrogels could also be one of the key factors playing a role in increased stiffness in these hydrogels. Together with previous reports showing the influence of fiber curvature amplitude and wavelength on fibroblast migration and polarization [42], investigating the influence of fibrotic ECM curvature on fibroblasts might reveal new insights into how fibroblast-ECM interactions are regulated by the physical state of the matrix structure. The stress relaxation behavior of the empty and fibroblast-encapsulated hydrogels showed that the changes in high density matrix and stiffness do not strongly influence the stress relaxation capacity of the fibers. It would be of interest to further investigate the altered stress relaxation of the fibers with respect to activation of the fibroblasts, as previously pre-stress in ECM fibers has been shown to release stored TGF-β from the ECM [43, 44]. Regardless, a recent report indicates that a microenvironment with slow stress capacity hinders cellular migration of mesenchymal spheroids [45]. Understanding how cellular migration plays a role in the profibrotic activation of fibroblasts with respect to organization of ECM requires further studies.

While our study establishes the interplay between the native microenvironment and the fibroblast-driven ECM remodeling responses in 3D, it has a few limitations. Human lung ECM (both control and IPF) hydrogels were variable between experimental runs. Although we generated a combined batch of ECM derived from 7 different donors to minimize this variation, sample-to-sample variation was still present in our results. Empty hydrogels harvested at each of the assessment time points also reflected this variation. However, we accounted for this variation during our analyses and only compared empty hydrogels to fibroblast-encapsulated hydrogels within the same time point and the same experimental run. Even with this comparison, it is not possible to rule out the fact that spontaneous fiber reorganization still continues during the course of 14 day cell culture in empty and fibroblast-encapsulated hydrogels. Another potential limitation of the study is related to collagen detection: as the starting weights of control and IPF lung ECM powders were the same, and as the majority of the remaining proteins within the dECMs were collagens, detecting small changes that might have been induced by the fibroblasts may not have been possible with the methodologies used in this study. While investigating changes in different collagens individually and also other ECM proteins was outside of the scope of this project, it is not possible to rule out that specific collagens may have been altered by the fibroblasts more than others, or that other ECM proteins including proteoglycans may have been involved. Lastly, viscoelastic stress relaxation of the control and IPF lung ECM-derived hydrogels is difficult, if at all possible, to resemble native tissue (a recognized limitation of this model [12, 32]), as opposed to the stiffness values that do recapitulate the patterns seen in lung tissues.

The model described in this study also provides opportunities for further research. Although our current model is based on fibroblast-ECM interactions, introducing other cell types such as epithelial cells, circulating immune cells or other mesenchymal cells would help mimicking the complex interplay between these cells and ECM during IPF or other fibrotic lung diseases. Moreover, this model system can greatly advance investigating cell responses following treatment with Nintedanib or Pirfenidone, which were initially performed on cell-derived matrices [46]. While our study focused on IPF, it has important implications for understanding the interplay between fibroblast-driven ECM-remodeling responses of fibroblasts and the fibrotic environment in many other fibrotic diseases and even cancer, reflecting the remodeled ECM associated with tumors. Understanding how activation of fibroblasts occurs in a fibrotic microenvironment has the potential to reveal additional intervention possibilities for the treatment of diseases involving fibrotic responses. Future studies utilizing this model could investigate if the fibroblast ECM remodeling responses differ in the presence of antifibrotic treatments that are currently approved for pulmonary fibrosis.

In summary, we examined how both primary lung control and IPF fibroblasts in a 3D fibrotic microenvironment interact with this environment and subsequently remodel their environment. Through employing native ECM from control and IPF lungs, most biochemical and biomechanical properties of control and IPF lungs were mimicked in these hydrogels, thereby presenting an innovative model system for investigating the interplay between the microenvironment and fibroblasts during the fibrotic process. Considering the lack of physiologically replicative models available for basic and translational research for generating treatment strategies targeting fibrosis, employing *in vitro* models derived from human-sourced materials can pave the way towards better understanding of fibrosis and potential drug discovery processes.

## Acknowledgments

Authors thank Mr. Albano Tosato for assistance in preparation of visuals.

## Data and materials availability

Estimates used to reach to reach the presented conclusions in this manuscript are included within the manuscript and/or added in the Supplementary Materials (*Supplementary Figures 7-42* for graphs with individual estimate points and statistical analyses of these individual points in *Supplementary Tables 1-36* are included in *Supplementary Document 3*). The raw data that support the findings of this study are available from the corresponding author upon reasonable request.

