## Supplementary Figures 1-6 for "Three dimensional fibrotic extracellular matrix directs microenvironment fiber remodeling by fibroblasts"

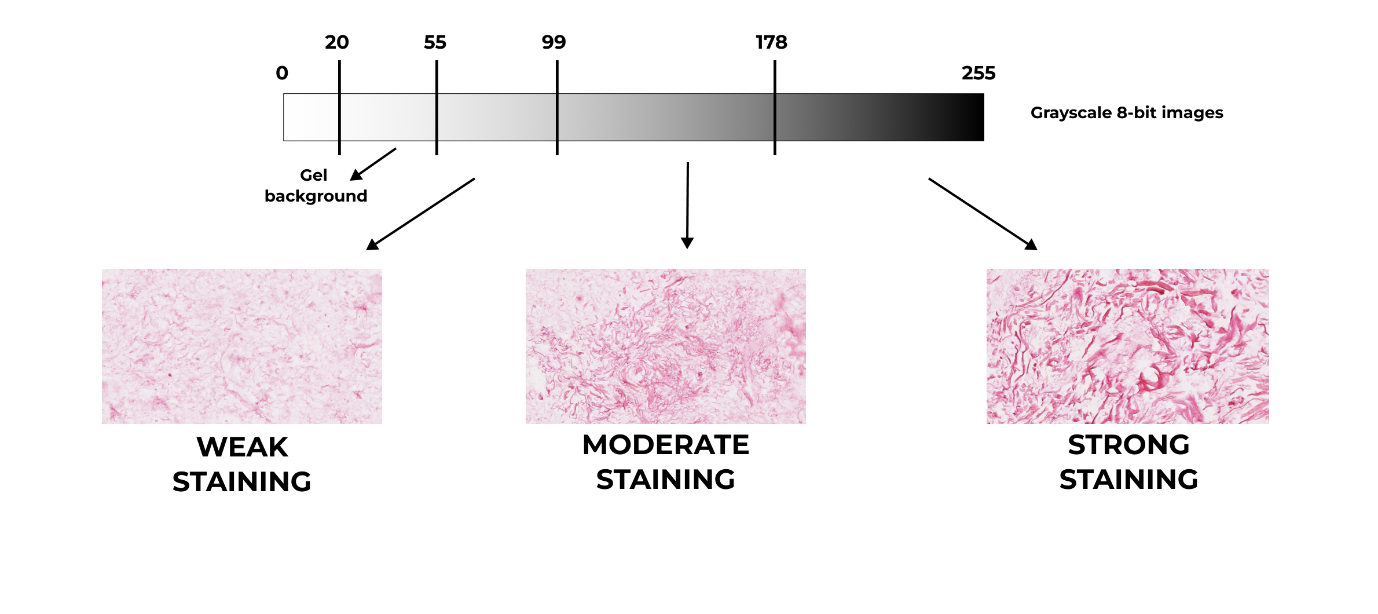

Supplementary Figure 1: Image analysis strategy used to analyze images of PicroSirius Red stained-slides. 8-bit images were converted to grayscale images and each pixel was assigned a strength between 0-255. Based on the strength values, the gel background was detected between 20 - 55 pixel strengths, and the remaining values (55 - 255) were divided into three categories to compare: weak staining (left), moderate staining (middle) and strong staining (right).

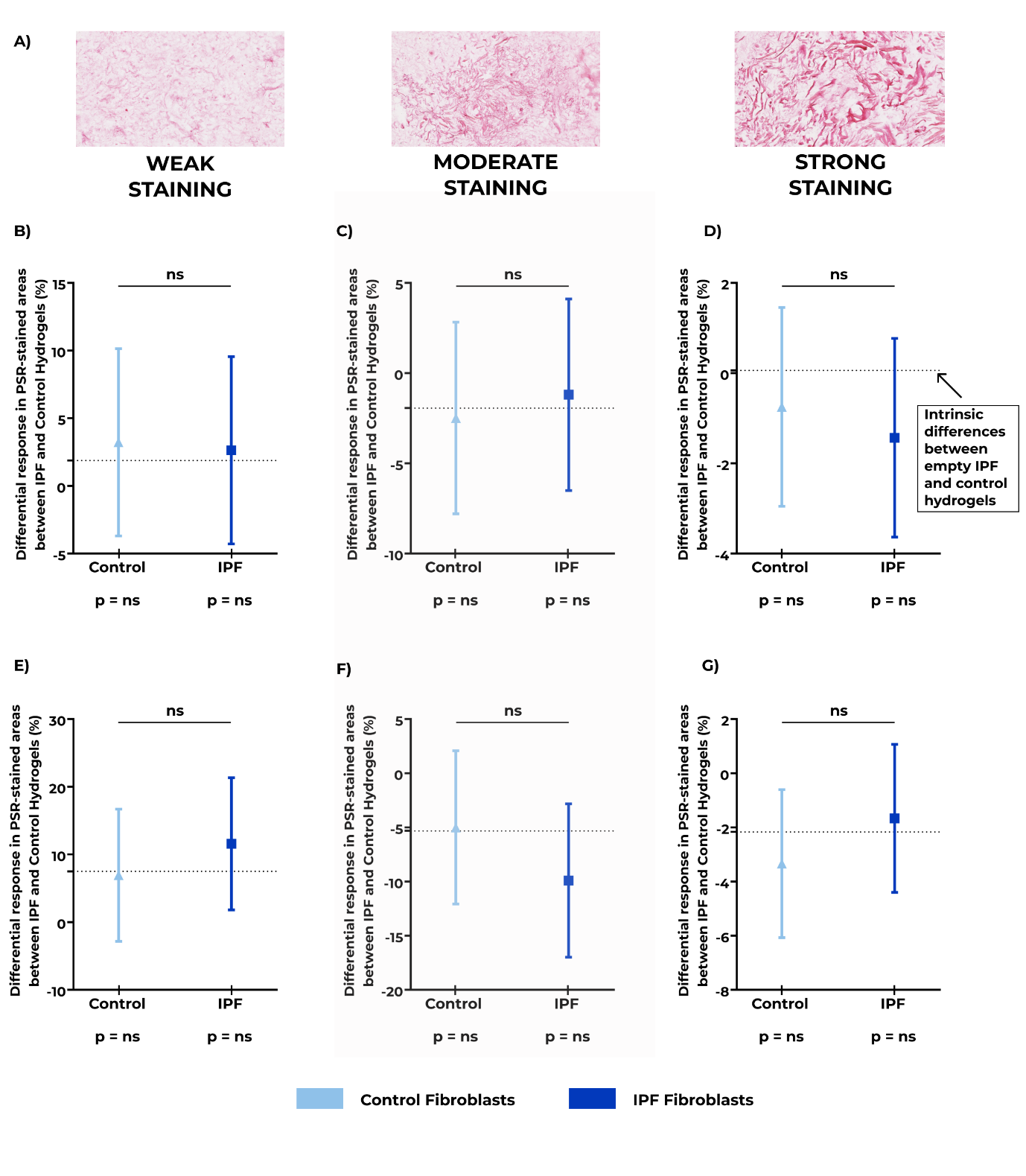

**Supplementary Figure 2:** Deposited collagen amounts in empty and fibroblast-encapsulated lung ECM-derived hydrogels. Control and IPF primary lung fibroblasts were encapsulated in control or IPF lung ECM-derived hydrogels and cultured for 7 or 14 days. Collagens were visualized using PicroSirius Red (PSR) staining on sections of paraffin-embedded fibroblast-seeded hydrogels and staining levels were compared with their corresponding empty hydrogel samples. A) Example images for the three levels of the strength of PSR staining. Day 7 responses with respect to B) weak C) moderate and D) strong PSR-stained areas (% area). Day 14 responses with respect to E) weak F) moderate and G) strong PSR-stained areas (% area). The dotted line shows the intrinsic difference between the empty IPF and control hydrogels. The estimate (± 95% confidence interval) shows the difference between the IPF and control hydrogels laden with control (light blue, triangle) or IPF (dark blue, square) fibroblasts. P values below each fibroblast group represent the differences induced by fibroblasts in IPF versus control hydrogels compared to the intrinsic difference between IPF and control empty hydrogels. P-values above the estimates indicate the differences between the responses of IPF and control fibroblasts in the different hydrogels. Applied statistical test: mixed-model analysis. ns: not significant, IPF: Idiopathic pulmonary fibrosis. n=6 for fibroblast donors, 1 image per sample was analyzed.

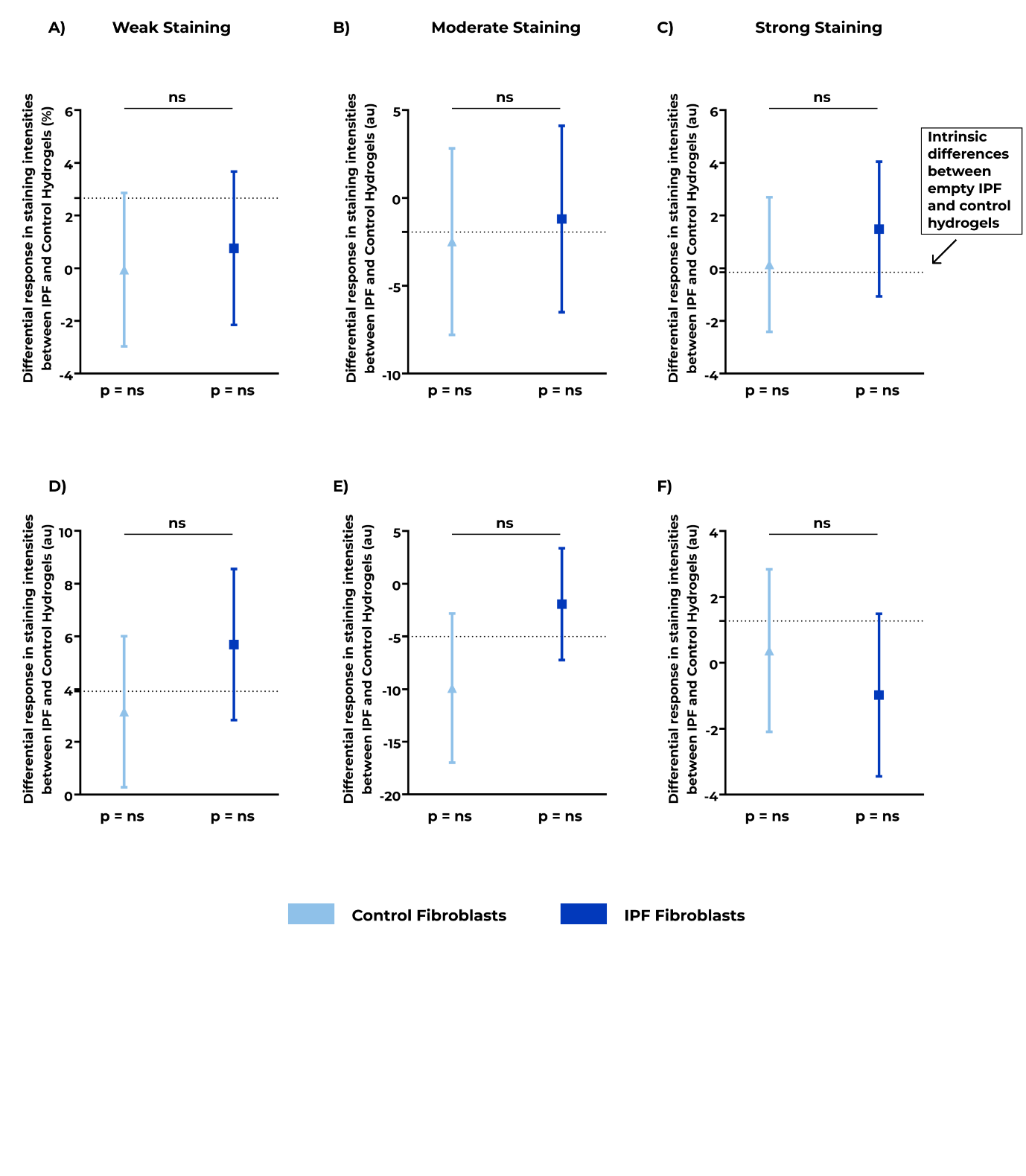
 **Supplementary Figure 3:** Deposited collagen intensities in empty and fibroblast-encapsulated lung ECM-derived hydrogels. Control and IPF primary lung fibroblasts were encapsulated in control or IPF lung ECM-derived hydrogels and cultured for 7 or 14 days. Collagens were visualized using PicroSirius Red (PSR) staining on sections of paraffin-embedded fibroblast-seeded hydrogels and staining intensities were compared with their corresponding empty hydrogel samples. Day 7 responses with respect to A) weak B) moderate and C) strong PSR-stained intensities (arbitrary unit (au)). Day 14 responses with respect to D) weak E) moderate and F) strong PSR-stained areas (au). The dotted line shows the intrinsic difference between the empty IPF and control hydrogels. The estimate (± 95% confidence interval) shows the difference between the IPF and control hydrogels laden with control (light blue, triangle) or IPF (dark blue, square) fibroblasts. P values below each fibroblast group represent the differences induced by fibroblasts in IPF versus control hydrogels compared to the intrinsic difference between IPF and control empty hydrogels. P-values above the estimates indicate the differences between the responses of IPF and control fibroblasts in the different hydrogels. Applied statistical test: mixed-model analysis. ns: not significant, IPF: Idiopathic pulmonary fibrosis. n=6 for fibroblast donors, 1 image per sample was analyzed.

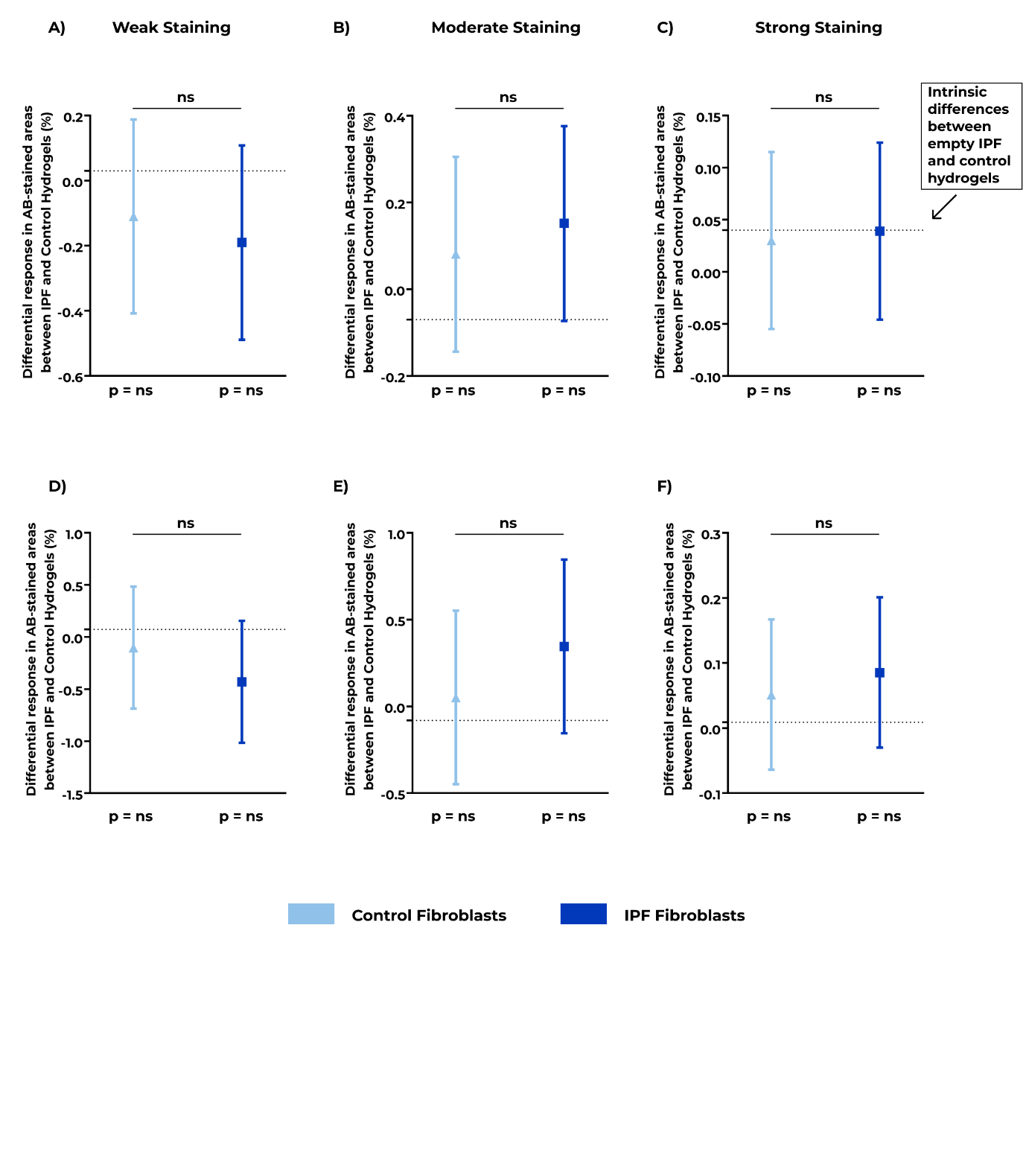
 **Supplementary Figure 4:** Deposited glycosaminoglycan amounts in empty and fibroblast-encapsulated lung ECM-derived hydrogels. Control and IPF primary lung fibroblasts were encapsulated in control or IPF lung ECM-derived hydrogels and cultured for 7 or 14 days. Glycosaminoglycans were visualized using Alcian Blue (AB) staining on sections of paraffin-embedded fibroblast-seeded hydrogels and staining levels were compared with their corresponding empty hydrogel samples. Day 7 responses with respect to A) weak B) moderate and C) strong AB-stained areas (% area). Day 14 responses with respect to D) weak E) moderate and F) strong AB-stained areas (% area). The dotted line shows the intrinsic difference between the empty IPF and control hydrogels. The estimate (± 95% confidence interval) shows the difference between the IPF and control hydrogels laden with control (light blue, triangle) or IPF (dark blue, square) fibroblasts. P values below each fibroblast group represent the differences induced by fibroblasts in IPF versus control hydrogels compared to the intrinsic difference between IPF and control empty hydrogels. P-values above the estimates indicate the differences between the responses of IPF and control fibroblasts in the different hydrogels. Applied statistical test: mixed-model analysis. ns: not significant, IPF: Idiopathic pulmonary fibrosis. n=6 for fibroblast donors, 1 image per sample was analyzed.

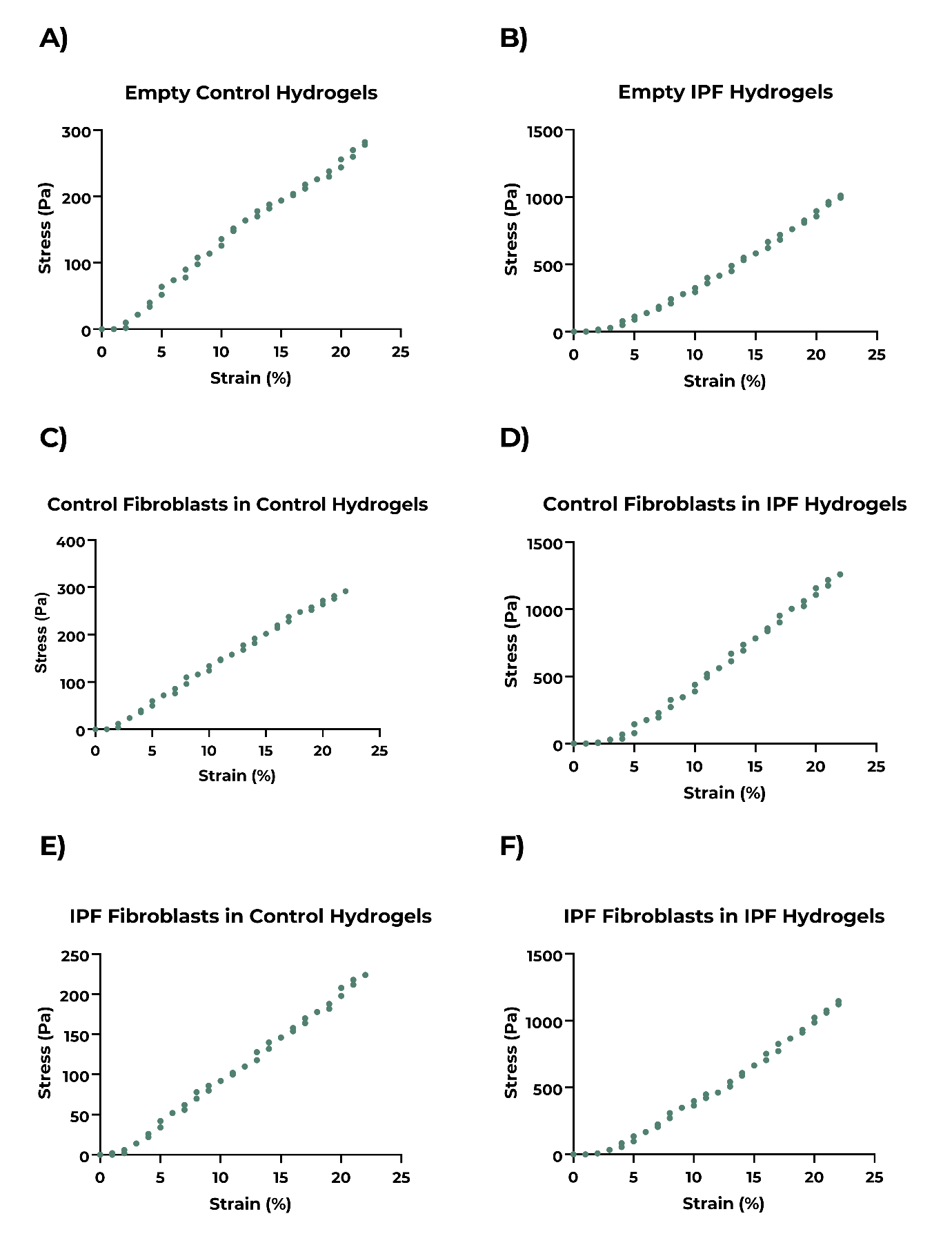
 **Supplementary Figure 5:** Representative stress-strain curves for the empty and fibroblast-encapsulated control and IPF hydrogels. A) Empty control hydrogels, B) Empty IPF hydrogels, C) Control fibroblasts in control hydrogels, D) Control fibroblasts in IPF hydrogels, E) IPF hydrogels in control hydrogels, F) IPF fibroblasts in IPF hydrogels.

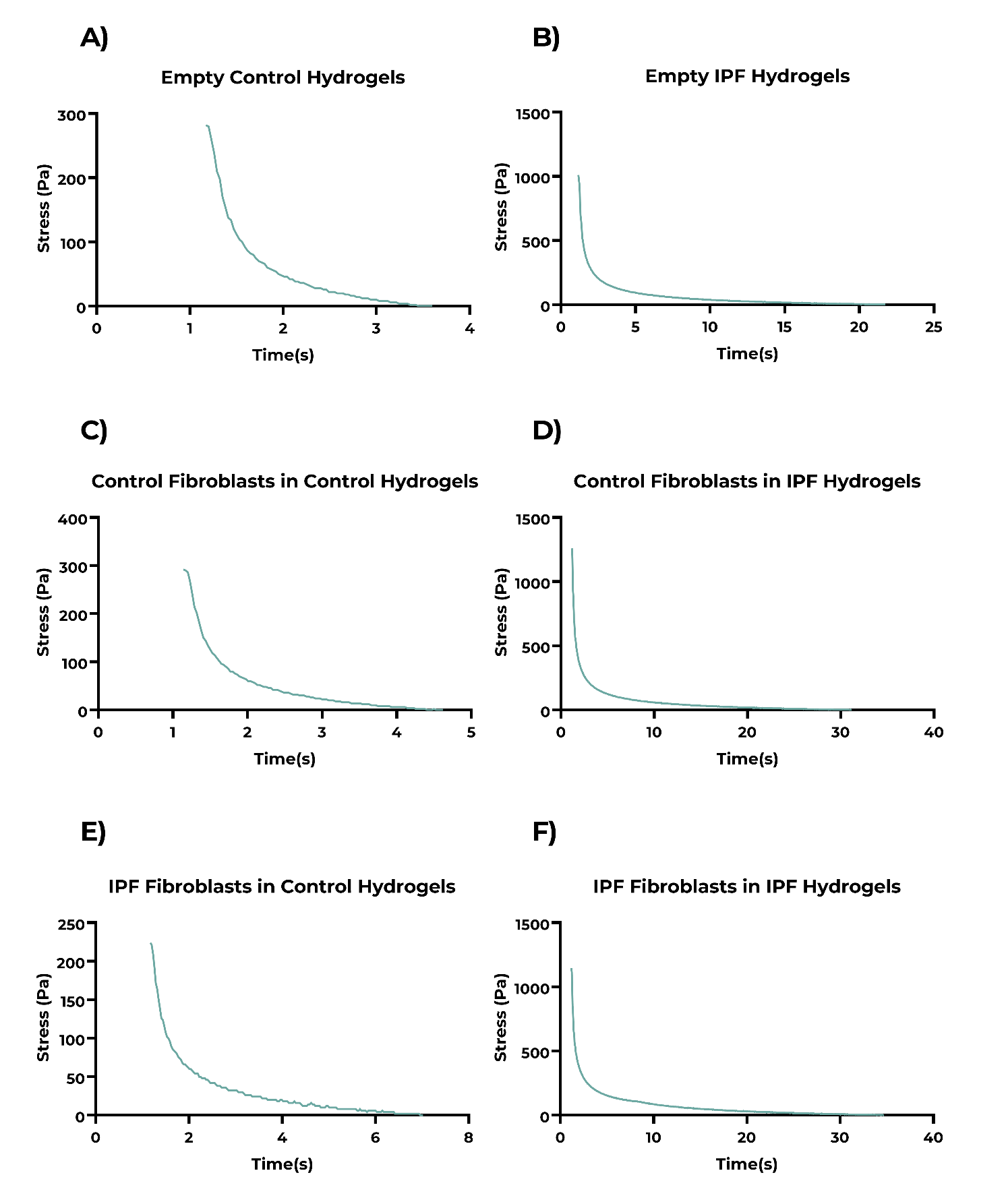

**Supplementary Figure 6:** Representative stress relaxation profiles for the empty and fibroblast-encapsulated control and IPF hydrogels. A) Empty control hydrogels, B) Empty IPF hydrogels, C) Control fibroblasts in control hydrogels, D) Control fibroblasts in IPF hydrogels, E) IPF hydrogels in control hydrogels, F) IPF fibroblasts in IPF hydrogels.
