## Supplementary Document 1 for "Three dimensional fibrotic extracellular matrix directs microenvironment fiber remodeling by fibroblasts"

**Supplementary Document 1:** ImageJ Macro used to analyze the images of PicroSirius Red-stained slides

name=getTitle();

print(name);

display=getTitle();

run("Set Measurements...", "area mean modal integrated median skewness kurtosis area_fraction limit display redirect=None decimal=6");

run("Colour Deconvolution", "vectors=[ Picro_Sirius_Red]");

// Vector Picro_Sirius_Red =

// Picro_Sirius_Red, 0.650,0.700,0.450, 0.150,0.750,0.400, 0.200,0.500,0.800

selectWindow(name+"-(Colour_1)");

setAutoThreshold("Default");

setThreshold(195, 200);

run("Measure");

setThreshold(156, 194);

run("Measure");

setThreshold(117, 155);

run("Measure");

setThreshold(78, 116);

run("Measure");

setThreshold(39, 77);

run("Measure");

setThreshold(0, 38);

run("Measure");

run("Close All");
