## Supplementary Document 2 for "Three dimensional fibrotic extracellular matrix directs microenvironment fiber remodeling by fibroblasts"

**Supplementary Document 2:** ImageJ Macro used to analyze the images of Alcian blue-stained slides

name=getTitle();

print(name);

display=getTitle();

run("Set Measurements...", "area mean modal integrated median skewness kurtosis area_fraction limit display redirect=None decimal=6");

run("Duplicate...", " ");

run("8-bit");

name1=getTitle();

rename(name1+"-(Colour_0)");

setAutoThreshold("Default");

setThreshold(0, 220);

run("Measure");

close();

run("Colour Deconvolution", "vectors=[ AlcianBlue]");

// AlcianBlue vector: AlcianBlue, 0.300,0.850,0.400, 0.800,0.550,0.250, 0.00000000,0.00000000,0.0000000

// Fill in below the thresholds for Colour2 image

selectWindow(name+"-(Colour_2)");

setAutoThreshold("Default");

setThreshold(0, 220);

run("Measure");

setThreshold(188, 220);

run("Measure");

setThreshold(150, 187);

run("Measure");

setThreshold(113, 149);

run("Measure");

setThreshold(75, 112);

run("Measure");

setThreshold(38, 74);

run("Measure");

setThreshold(0, 37);

run("Measure");

run("Close All");
