## Supplementary Document 3 for "Three dimensional fibrotic extracellular matrix directs microenvironment fiber remodeling by fibroblasts"

| Supplementary Table 1: Individual comparisons between the different groups of empty and fibroblast-encapsulated control and IPF hydrogels for weakly stained PicroSirius Red area on day 7 samples | | | | | | | |
| --- | --- | --- | --- | --- | --- | --- | --- |
|  |  | Control Hydrogels | | | IPF Hydrogels | | |
|  |  | Empty | Control Fb | IPF Fb | Empty | Control Fb | IPF Fb |
| Control Hydrogels | Empty |  | 0.478 | 0.364 | 0.583 |  |  |
|  | Control Fb | 0.478 |  | 0.111 |  |  |  |
|  | IPF Fb | 0.364 | 0.111 |  |  |  |  |
| IPF Hydrogels | Empty | 0.583 |  |  |  | 0.273 | 0.489 |
|  | Control Fb |  |  |  | 0.273 |  | 0.079 |
|  | IPF Fb |  |  |  | 0.489 | 0.079 |  |
|  |  |  |  | Not significant | |  |  |
|  |  |  |  | Significant | |  |  |
|  |  |  |  | Comparison not applicable | |  |  |

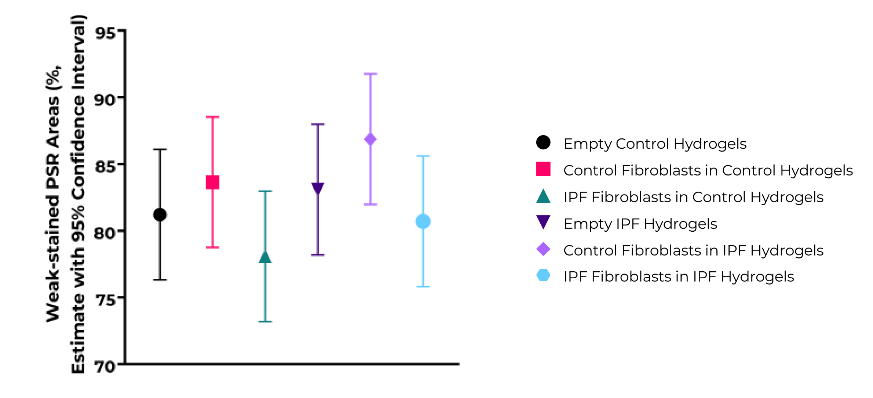

Supplementary Figure 7: Individual comparisons between the different groups of empty and fibroblast-encapsulated control and IPF hydrogels for the estimates of weakly stained PicroSirius Red (PSR) area (%) for day 7 samples.

| Supplementary Table 2: Individual comparisons between the different groups of empty and fibroblast-encapsulated control and IPF hydrogels for weakly stained PicroSirius Red area on day 14 samples | | | | | | | |
| --- | --- | --- | --- | --- | --- | --- | --- |
|  |  | Control Hydrogels | | | IPF Hydrogels | | |
|  |  | Empty | Control Fb | IPF Fb | Empty | Control Fb | IPF Fb |
| Control Hydrogels | Empty |  | 0.371 | 0.941 | 0.126 |  |  |
|  | Control Fb | 0.371 |  | 0.334 |  |  |  |
|  | IPF Fb | 0.941 | 0.334 |  |  |  |  |
| IPF Hydrogels | Empty | 0.126 |  |  |  | 0.435 | 0.439 |
|  | Control Fb |  |  |  | 0.435 |  | 0.995 |
|  | IPF Fb |  |  |  | 0.439 | 0.995 |  |
|  |  |  |  | Not significant | |  |  |
|  |  |  |  | Significant | |  |  |
|  |  |  |  | Comparison not applicable | |  |  |

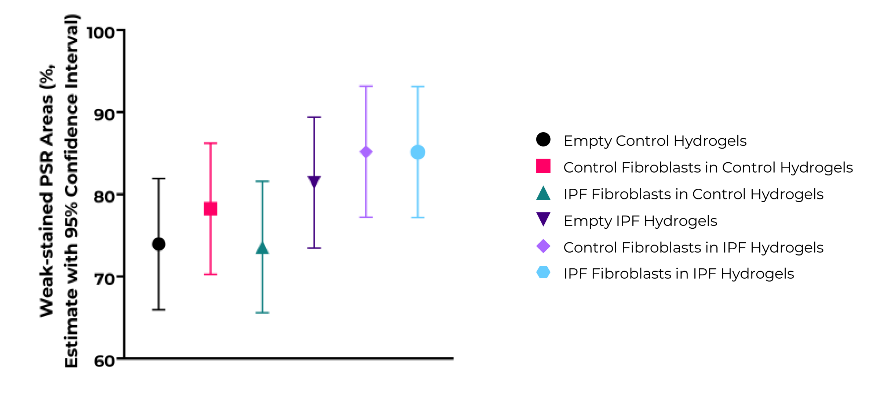

Supplementary Figure 8: Individual comparisons between the different groups of empty and fibroblast-encapsulated control and IPF hydrogels for the estimates of weakly stained PicroSirius Red (PSR) area (%) for day 14 samples.

| Supplementary Table 3: Individual comparisons between the different groups of empty and fibroblast-encapsulated control and IPF hydrogels for moderately stained PicroSirius Red area on day 7 samples | | | | | | | |
| --- | --- | --- | --- | --- | --- | --- | --- |
|  |  | Control Hydrogels | | | IPF Hydrogels | | |
|  |  | Empty | Control Fb | IPF Fb | Empty | Control Fb | IPF Fb |
| Control Hydrogels | Empty |  | 0.465 | 0.398 | 0.462 |  |  |
|  | Control Fb | 0.639 |  | 0.121 |  |  |  |
|  | IPF Fb | 0.412 | 0.201 |  |  |  |  |
| IPF Hydrogels | Empty | 0.955 |  |  |  | 0.350 | 0.263 |
|  | Control Fb |  |  |  | 0.230 |  | 0.045 |
|  | IPF Fb |  |  |  | 0.584 | 0.045 |  |
|  |  |  |  | Not significant | |  |  |
|  |  |  |  | Significant | |  |  |
|  |  |  |  | Comparison not applicable | |  |  |

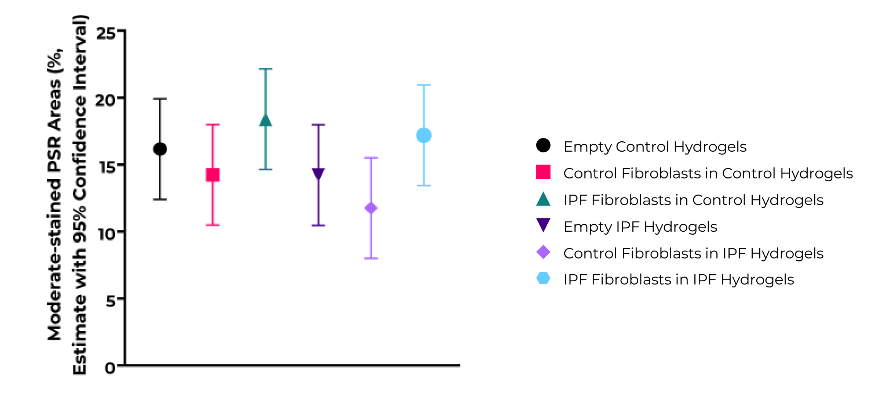

Supplementary Figure 9: Individual comparisons between the different groups of empty and fibroblast-encapsulated control and IPF hydrogels for the estimates of moderately stained PicroSirius Red (PSR) area (%) for day 7 samples.

| Supplementary Table 4: Individual comparisons between the different groups of empty and fibroblast-encapsulated control and IPF hydrogels for moderately stained PicroSirius Red area on day 14 samples | | | | | | | |
| --- | --- | --- | --- | --- | --- | --- | --- |
|  |  | Control Hydrogels | | | IPF Hydrogels | | |
|  |  | Empty | Control Fb | IPF Fb | Empty | Control Fb | IPF Fb |
| Control Hydrogels | Empty |  | 0.405 | 0.638 | 0.134 |  |  |
|  | Control Fb | 0.405 |  | 0.198 |  |  |  |
|  | IPF Fb | 0.638 | 0.198 |  |  |  |  |
| IPF Hydrogels | Empty | 0.134 |  |  |  | 0.458 | 0.399 |
|  | Control Fb |  |  |  | 0.458 |  | 0.919 |
|  | IPF Fb |  |  |  | 0.399 | 0.919 |  |
|  |  |  |  | Not significant | |  |  |
|  |  |  |  | Significant | |  |  |
|  |  |  |  | Comparison not applicable | |  |  |

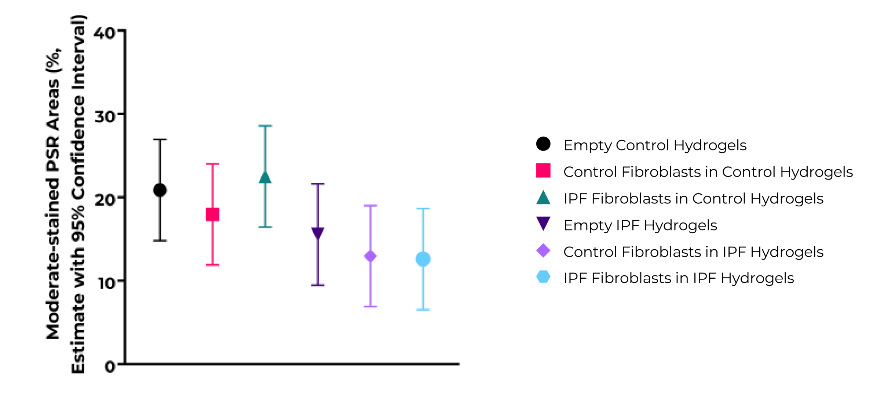

Supplementary Figure 10: Individual comparisons between the different groups of empty and fibroblast-encapsulated control and IPF hydrogels for the estimates of moderately stained PicroSirius Red (PSR) area (%) for day 14 samples.

| Supplementary Table 5: Individual comparisons between the different groups of empty and fibroblast-encapsulated control and IPF hydrogels for strongly stained PicroSirius Red area on day 7 samples | | | | | | | |
| --- | --- | --- | --- | --- | --- | --- | --- |
|  |  | Control Hydrogels | | | IPF Hydrogels | | |
|  |  | Empty | Control Fb | IPF Fb | Empty | Control Fb | IPF Fb |
| Control Hydrogels | Empty |  | 0.639 | 0.412 | 0.955 |  |  |
|  | Control Fb | 0.639 |  | 0.201 |  |  |  |
|  | IPF Fb | 0.412 | 0.201 |  |  |  |  |
| IPF Hydrogels | Empty | 0.955 |  |  |  | 0.230 | 0.584 |
|  | Control Fb |  |  |  | 0.230 |  | 0.507 |
|  | IPF Fb |  |  |  | 0.584 | 0.507 |  |
|  |  |  |  | Not significant | |  |  |
|  |  |  |  | Significant | |  |  |
|  |  |  |  | Comparison not applicable | |  |  |

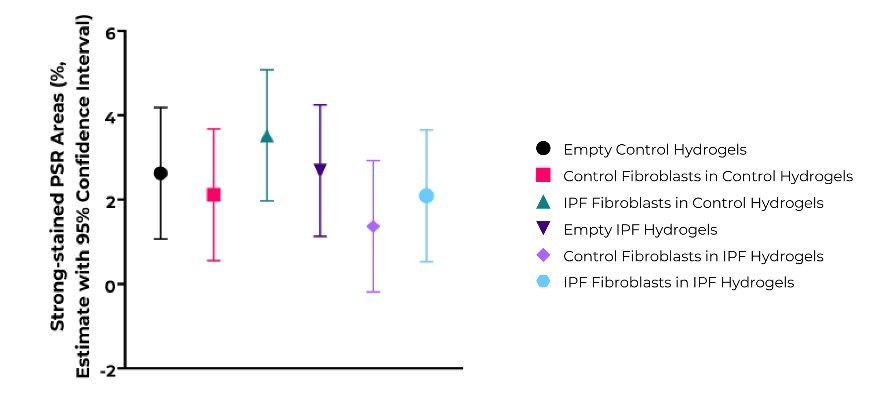

Supplementary Figure 11: Individual comparisons between the different groups of empty and fibroblast-encapsulated control and IPF hydrogels for the estimates of strongly stained PicroSirius Red (PSR) area (%) for day 7 samples.

| Supplementary Table 6: Individual comparisons between the different groups of empty and fibroblast-encapsulated control and IPF hydrogels for strongly stained PicroSirius Red area on day 14 samples | | | | | | | |
| --- | --- | --- | --- | --- | --- | --- | --- |
|  |  | Control Hydrogels | | | IPF Hydrogels | | |
|  |  | Empty | Control Fb | IPF Fb | Empty | Control Fb | IPF Fb |
| Control Hydrogels | Empty |  | 1.000 | 0.343 | 0.115 |  |  |
|  | Control Fb | 1.000 |  | 0.343 |  |  |  |
|  | IPF Fb | 0.343 | 0.343 |  |  |  |  |
| IPF Hydrogels | Empty | 0.115 |  |  |  | 0.387 | 0.563 |
|  | Control Fb |  |  |  | 0.387 |  | 0.771 |
|  | IPF Fb |  |  |  | 0.563 | 0.771 |  |
|  |  |  |  | Not significant | |  |  |
|  |  |  |  | Significant | |  |  |
|  |  |  |  | Comparison not applicable | |  |  |

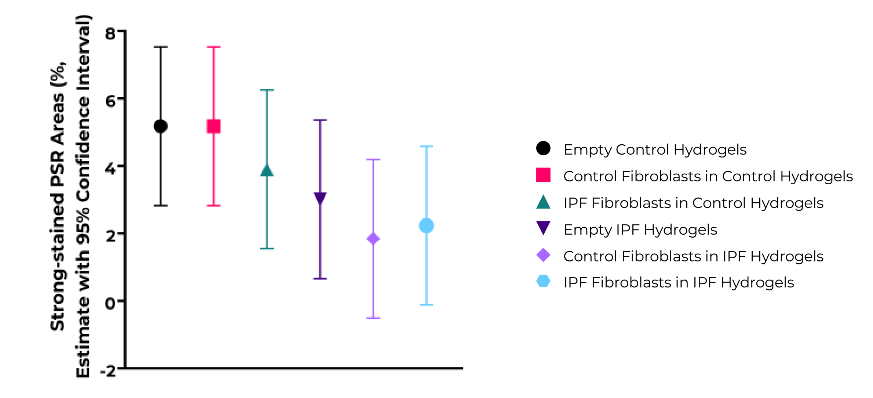

Supplementary Figure 12: Individual comparisons between the different groups of empty and fibroblast-encapsulated control and IPF hydrogels for the estimates of strongly stained PicroSirius Red (PSR) area (%) for day 14 samples.

| Supplementary Table 7: Individual comparisons between the different groups of empty and fibroblast-encapsulated control and IPF hydrogels for mean intensity of weakly stained PicroSirius Red on day 7 samples | | | | | | | |
| --- | --- | --- | --- | --- | --- | --- | --- |
|  |  | Control Hydrogels | | | IPF Hydrogels | | |
|  |  | Empty | Control Fb | IPF Fb | Empty | Control Fb | IPF Fb |
| Control Hydrogels | Empty |  | 0.600 | 0.663 | 0.071 |  |  |
|  | Control Fb | 0.600 |  | 0.181 |  |  |  |
|  | IPF Fb | 0.663 | 0.181 |  |  |  |  |
| IPF Hydrogels | Empty | 0.071 |  |  |  | 0.682 | 0.026 |
|  | Control Fb |  |  |  | 0.682 |  | 0.062 |
|  | IPF Fb |  |  |  | 0.026 | 0.062 |  |
|  |  |  |  | Not significant | |  |  |
|  |  |  |  | Significant | |  |  |
|  |  |  |  | Comparison not applicable | |  |  |

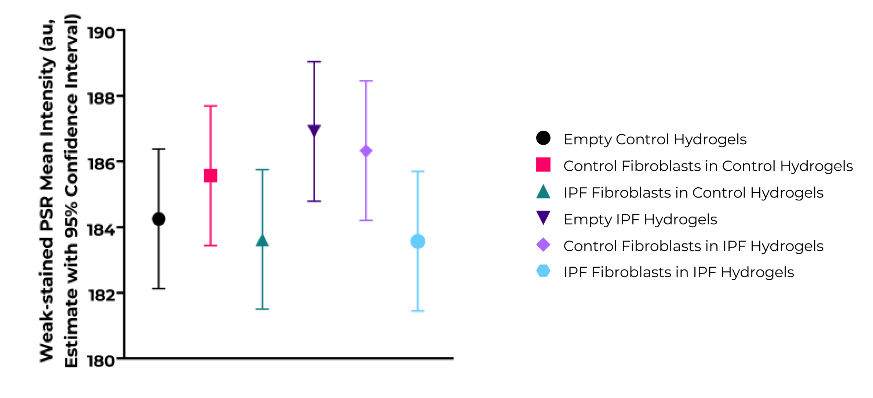

Supplementary Figure 13: Individual comparisons between the different groups of empty and fibroblast-encapsulated control and IPF hydrogels for the estimates of mean intensity of weakly stained PicroSirius Red (PSR) area (au) for day 7 samples.

| Supplementary Table 8: Individual comparisons between the different groups of empty and fibroblast-encapsulated control and IPF hydrogels for mean intensity of weakly stained PicroSirius Red on day 14 samples | | | | | | | |
| --- | --- | --- | --- | --- | --- | --- | --- |
|  |  | Control Hydrogels | | | IPF Hydrogels | | |
|  |  | Empty | Control Fb | IPF Fb | Empty | Control Fb | IPF Fb |
| Control Hydrogels | Empty |  | 0.403 | 0.057 | 0.009 |  |  |
|  | Control Fb | 0.403 |  | 0.165 |  |  |  |
|  | IPF Fb | 0.057 | 0.165 |  |  |  |  |
| IPF Hydrogels | Empty | 0.009 |  |  |  | 0.720 | 0.493 |
|  | Control Fb |  |  |  | 0.720 |  | 0.691 |
|  | IPF Fb |  |  |  | 0.493 | 0.691 |  |
|  |  |  |  | Not significant | |  |  |
|  |  |  |  | Significant | |  |  |
|  |  |  |  | Comparison not applicable | |  |  |

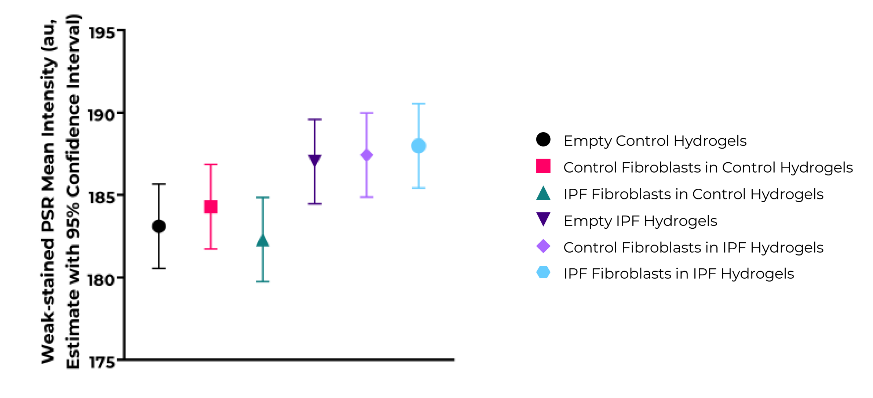

Supplementary Figure 14: Individual comparisons between the different groups of empty and fibroblast-encapsulated control and IPF hydrogels for the estimates of mean intensity of weakly stained PicroSirius Red (PSR) area (au) for day 14 samples.

| Supplementary Table 9: Individual comparisons between the different groups of empty and fibroblast-encapsulated control and IPF hydrogels for mean intensity of moderately stained PicroSirius Red on day 7 samples | | | | | | | |
| --- | --- | --- | --- | --- | --- | --- | --- |
|  |  | Control Hydrogels | | | IPF Hydrogels | | |
|  |  | Empty | Control Fb | IPF Fb | Empty | Control Fb | IPF Fb |
| Control Hydrogels | Empty |  | 0.465 | 0.398 | 0.462 |  |  |
|  | Control Fb | 0.465 |  | 0.121 |  |  |  |
|  | IPF Fb | 0.398 | 0.121 |  |  |  |  |
| IPF Hydrogels | Empty | 0.462 |  |  |  | 0.350 | 0.263 |
|  | Control Fb |  |  |  | 0.350 |  | 0.045 |
|  | IPF Fb |  |  |  | 0.263 | 0.045 |  |
|  |  |  |  | Not significant | |  |  |
|  |  |  |  | Significant | |  |  |
|  |  |  |  | Comparison not applicable | |  |  |

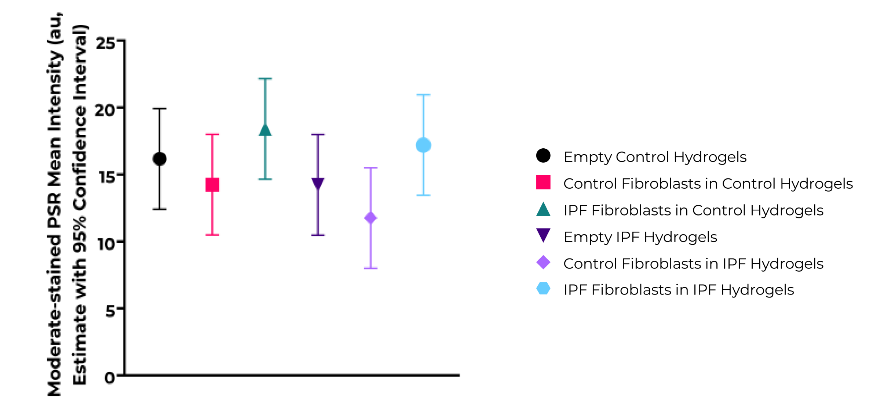

Supplementary Figure 15: Individual comparisons between the different groups of empty and fibroblast-encapsulated control and IPF hydrogels for the estimates of mean intensity of moderately stained PicroSirius Red (PSR) area (au) for day 7 samples.

| Supplementary Table 10: Individual comparisons between the different groups of empty and fibroblast-encapsulated control and IPF hydrogels for mean intensity of moderately stained PicroSirius Red on day 14 samples | | | | | | | |
| --- | --- | --- | --- | --- | --- | --- | --- |
|  |  | Control Hydrogels | | | IPF Hydrogels | | |
|  |  | Empty | Control Fb | IPF Fb | Empty | Control Fb | IPF Fb |
| Control Hydrogels | Empty |  | 0.405 | 0.198 | 0.158 |  |  |
|  | Control Fb | 0.405 |  | 0.198 |  |  |  |
|  | IPF Fb | 0.198 | 0.198 |  |  |  |  |
| IPF Hydrogels | Empty | 0.158 |  |  |  | 0.458 | 0.399 |
|  | Control Fb |  |  |  | 0.458 |  | 0.919 |
|  | IPF Fb |  |  |  | 0.399 | 0.919 |  |
|  |  |  |  | Not significant | |  |  |
|  |  |  |  | Significant | |  |  |
|  |  |  |  | Comparison not applicable | |  |  |

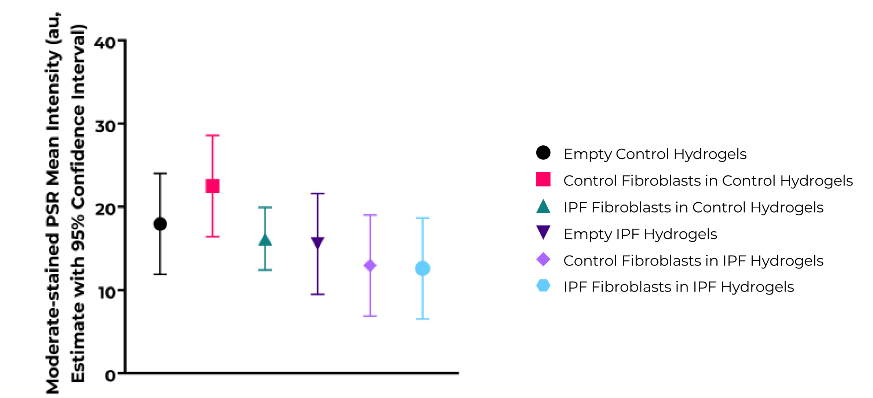

Supplementary Figure 16: Individual comparisons between the different groups of empty and fibroblast-encapsulated control and IPF hydrogels for the estimates of mean intensity of moderately stained PicroSirius Red (PSR) area (au) for day 14 samples.

| Supplementary Table 11: Individual comparisons between the different groups of empty and fibroblast-encapsulated control and IPF hydrogels for mean intensity of strongly stained PicroSirius Red on day 7 samples | | | | | | | |
| --- | --- | --- | --- | --- | --- | --- | --- |
|  |  | Control Hydrogels | | | IPF Hydrogels | | |
|  |  | Empty | Control Fb | IPF Fb | Empty | Control Fb | IPF Fb |
| Control Hydrogels | Empty |  | 0.702 | 0.431 | 0.903 |  |  |
|  | Control Fb | 0.702 |  | 0.246 |  |  |  |
|  | IPF Fb | 0.431 | 0.246 |  |  |  |  |
| IPF Hydrogels | Empty | 0.903 |  |  |  | 0.538 | 0.612 |
|  | Control Fb |  |  |  | 0.538 |  | 0.914 |
|  | IPF Fb |  |  |  | 0.612 | 0.914 |  |
|  |  |  |  | Not significant | |  |  |
|  |  |  |  | Significant | |  |  |
|  |  |  |  | Comparison not applicable | |  |  |

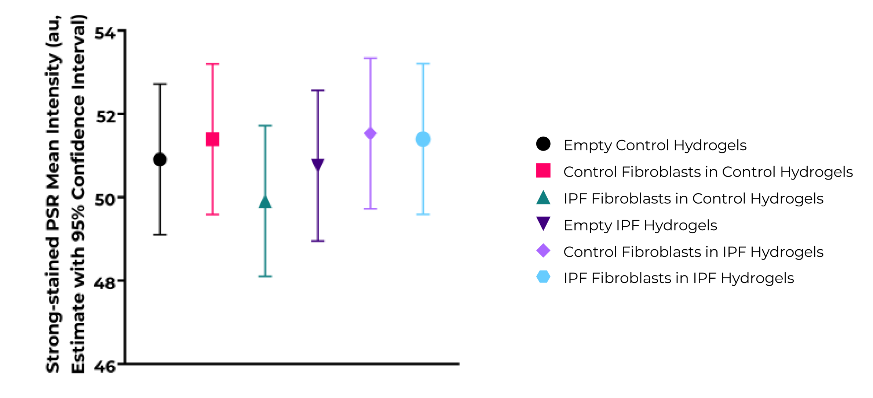

Supplementary Figure 17: Individual comparisons between the different groups of empty and fibroblast-encapsulated control and IPF hydrogels for the estimates of mean intensity of strongly stained PicroSirius Red (PSR) area (au) for day 7 samples.

| Supplementary Table 12: Individual comparisons between the different groups of empty and fibroblast-encapsulated control and IPF hydrogels for mean intensity of strongly stained PicroSirius Red on day 14 samples | | | | | | | |
| --- | --- | --- | --- | --- | --- | --- | --- |
|  |  | Control Hydrogels | | | IPF Hydrogels | | |
|  |  | Empty | Control Fb | IPF Fb | Empty | Control Fb | IPF Fb |
| Control Hydrogels | Empty |  | 0.701 | 0.296 | 0.298 |  |  |
|  | Control Fb | 0.701 |  | 0.503 |  |  |  |
|  | IPF Fb | 0.296 | 0.503 |  |  |  |  |
| IPF Hydrogels | Empty | 0.298 |  |  |  | 0.719 | 0.425 |
|  | Control Fb |  |  |  | 0.719 |  | 0.659 |
|  | IPF Fb |  |  |  | 0.425 | 0.659 |  |
|  |  |  |  | Not significant | |  |  |
|  |  |  |  | Significant | |  |  |
|  |  |  |  | Comparison not applicable | |  |  |

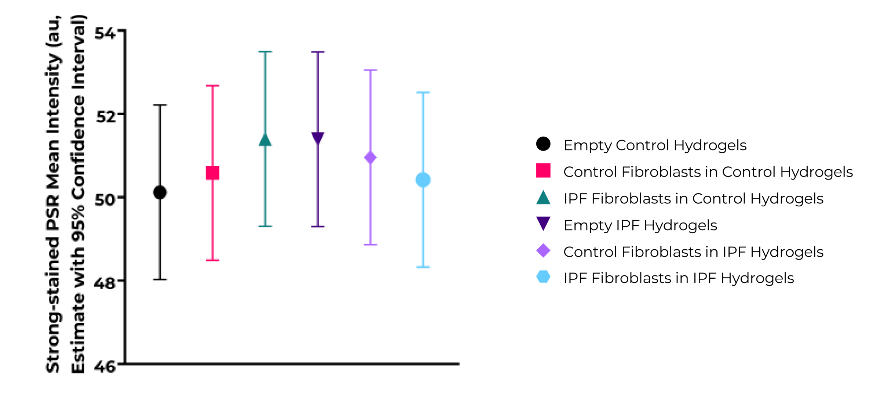

Supplementary Figure 18: Individual comparisons between the different groups of empty and fibroblast-encapsulated control and IPF hydrogels for the estimates of mean intensity of strongly stained PicroSirius Red (PSR) area (au) for day 14 samples.

| Supplementary Table 13: Individual comparisons between the different groups of empty and fibroblast-encapsulated control and IPF hydrogels for weakly stained Alcian Blue area on day 7 samples | | | | | | | |
| --- | --- | --- | --- | --- | --- | --- | --- |
|  |  | Control Hydrogels | | | IPF Hydrogels | | |
|  |  | Empty | Control Fb | IPF Fb | Empty | Control Fb | IPF Fb |
| Control Hydrogels | Empty |  | 0.281 | 0.080 | 0.696 |  |  |
|  | Control Fb | 0.281 |  | 0.453 |  |  |  |
|  | IPF Fb | 0.080 | 0.453 |  |  |  |  |
| IPF Hydrogels | Empty | 0.696 |  |  |  | 1.000 | 0.827 |
|  | Control Fb |  |  |  | 1.000 |  | 0.827 |
|  | IPF Fb |  |  |  | 0.827 | 0.827 |  |
|  |  |  |  | Not significant | |  |  |
|  |  |  |  | Significant | |  |  |
|  |  |  |  | Comparison not applicable | |  |  |

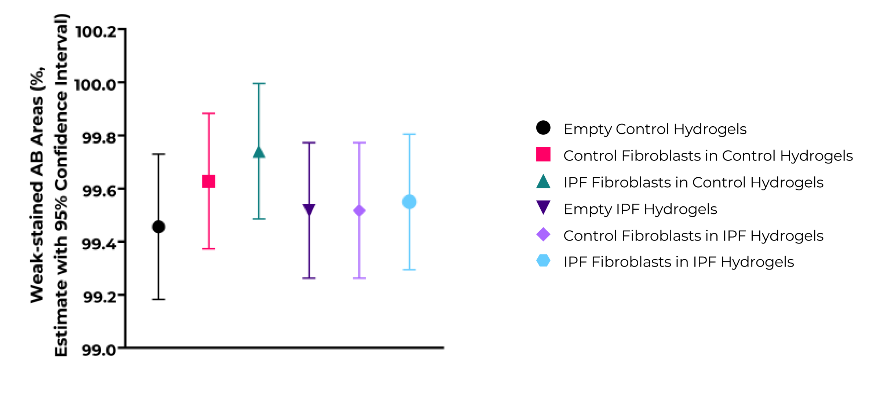

Supplementary Figure 19: Individual comparisons between the different groups of empty and fibroblast-encapsulated control and IPF hydrogels for the estimates of weakly stained Alcian blue (AB) area (%) for day 7 samples.

| Supplementary Table 14: Individual comparisons between the different groups of empty and fibroblast-encapsulated control and IPF hydrogels for weakly stained Alcian Blue area on day 14 samples | | | | | | | |
| --- | --- | --- | --- | --- | --- | --- | --- |
|  |  | Control Hydrogels | | | IPF Hydrogels | | |
|  |  | Empty | Control Fb | IPF Fb | Empty | Control Fb | IPF Fb |
| Control Hydrogels | Empty |  | 0.634 | 0.487 | 0.749 |  |  |
|  | Control Fb | 0.634 |  | 0.224 |  |  |  |
|  | IPF Fb | 0.487 | 0.224 |  |  |  |  |
| IPF Hydrogels | Empty | 0.749 |  |  |  | 0.241 | 0.289 |
|  | Control Fb |  |  |  | 0.241 |  | 0.908 |
|  | IPF Fb |  |  |  | 0.289 | 0.908 |  |
|  |  |  |  | Not significant | |  |  |
|  |  |  |  | Significant | |  |  |
|  |  |  |  | Comparison not applicable | |  |  |

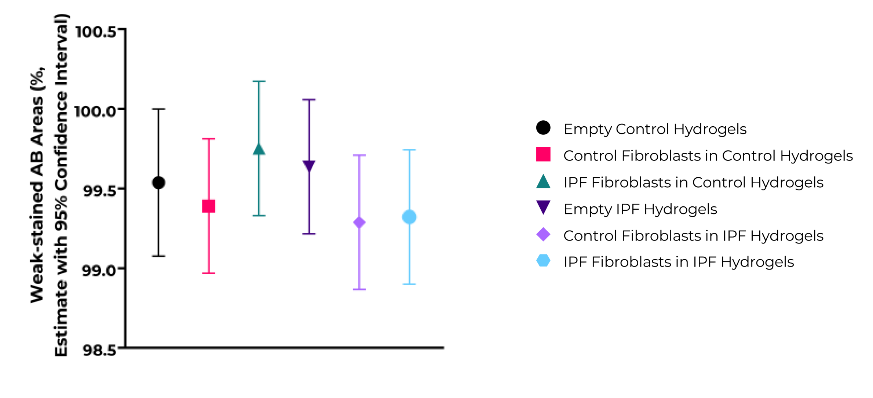

Supplementary Figure 20: Individual comparisons between the different groups of empty and fibroblast-encapsulated control and IPF hydrogels for the estimates of weakly stained Alcian blue (AB) area (%) for day 14 samples.

| Supplementary Table 15: Individual comparisons between the different groups of empty and fibroblast-encapsulated control and IPF hydrogels for moderately stained Alcian Blue area on day 7 samples | | | | | | | |
| --- | --- | --- | --- | --- | --- | --- | --- |
|  |  | Control Hydrogels | | | IPF Hydrogels | | |
|  |  | Empty | Control Fb | IPF Fb | Empty | Control Fb | IPF Fb |
| Control Hydrogels | Empty |  | 0.234 | 0.064 | 0.430 |  |  |
|  | Control Fb | 0.234 |  | 0.456 |  |  |  |
|  | IPF Fb | 0.064 | 0.456 |  |  |  |  |
| IPF Hydrogels | Empty | 0.430 |  |  |  | 0.780 | 0.871 |
|  | Control Fb |  |  |  | 0.780 |  | 0.906 |
|  | IPF Fb |  |  |  | 0.871 | 0.906 |  |
|  |  |  |  | Not significant | |  |  |
|  |  |  |  | Significant | |  |  |
|  |  |  |  | Comparison not applicable | |  |  |

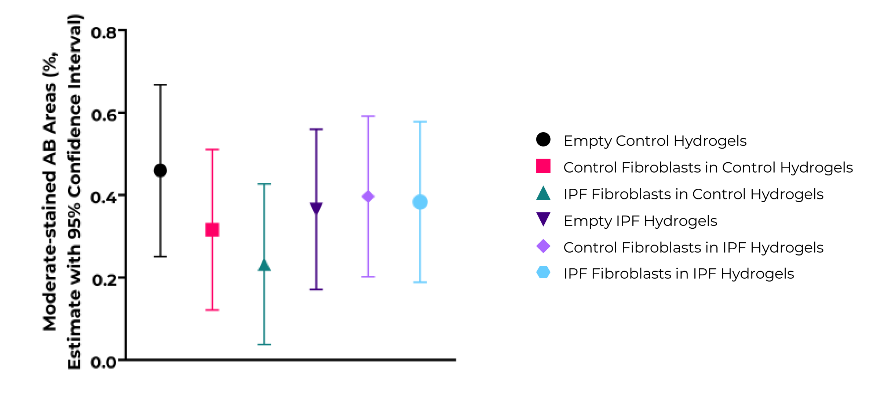

Supplementary Figure 21: Individual comparisons between the different groups of empty and fibroblast-encapsulated control and IPF hydrogels for the estimates of moderately stained Alcian blue (AB) area (%) for day 7 samples.

| Supplementary Table 16: Individual comparisons between the different groups of empty and fibroblast-encapsulated control and IPF hydrogels for moderately stained Alcian Blue area on day 14 samples | | | | | | | |
| --- | --- | --- | --- | --- | --- | --- | --- |
|  |  | Control Hydrogels | | | IPF Hydrogels | | |
|  |  | Empty | Control Fb | IPF Fb | Empty | Control Fb | IPF Fb |
| Control Hydrogels | Empty |  | 0.642 | 0.467 | 0.683 |  |  |
|  | Control Fb | 0.642 |  | 0.215 |  |  |  |
|  | IPF Fb | 0.467 | 0.215 |  |  |  |  |
| IPF Hydrogels | Empty | 0.683 |  |  |  | 0.267 | 0.303 |
|  | Control Fb |  |  |  | 0.267 |  | 0.934 |
|  | IPF Fb |  |  |  | 0.303 | 0.934 |  |
|  |  |  |  | Not significant | |  |  |
|  |  |  |  | Significant | |  |  |
|  |  |  |  | Comparison not applicable | |  |  |

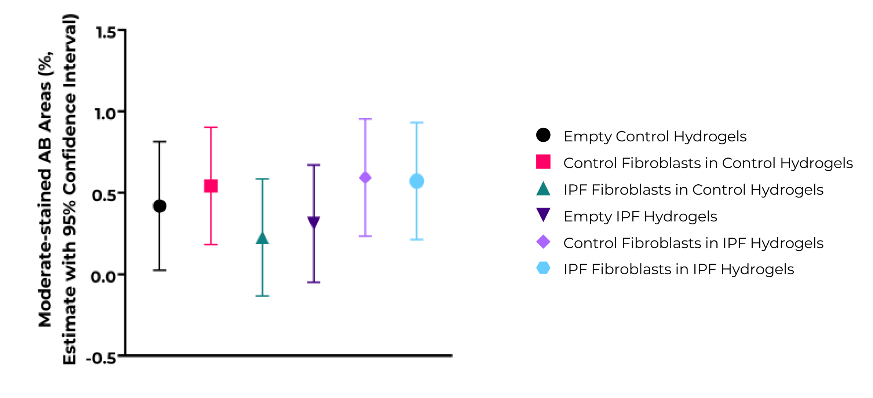

Supplementary Figure 22: Individual comparisons between the different groups of empty and fibroblast-encapsulated control and IPF hydrogels for the estimates of moderately stained Alcian blue (AB) area (%) for day 14 samples.

| Supplementary Table 17: Individual comparisons between the different groups of empty and fibroblast-encapsulated control and IPF hydrogels for strongly stained Alcian Blue area on day 7 samples | | | | | | | |
| --- | --- | --- | --- | --- | --- | --- | --- |
|  |  | Control Hydrogels | | | IPF Hydrogels | | |
|  |  | Empty | Control Fb | IPF Fb | Empty | Control Fb | IPF Fb |
| Control Hydrogels | Empty |  | 0.507 | 0.201 | 0.488 |  |  |
|  | Control Fb | 0.507 |  | 0.505 |  |  |  |
|  | IPF Fb | 0.201 | 0.505 |  |  |  |  |
| IPF Hydrogels | Empty | 0.488 |  |  |  | 0.463 | 0.240 |
|  | Control Fb |  |  |  | 0.463 |  | 0.650 |
|  | IPF Fb |  |  |  | 0.240 | 0.650 |  |
|  |  |  |  | Not significant | |  |  |
|  |  |  |  | Significant | |  |  |
|  |  |  |  | Comparison not applicable | |  |  |

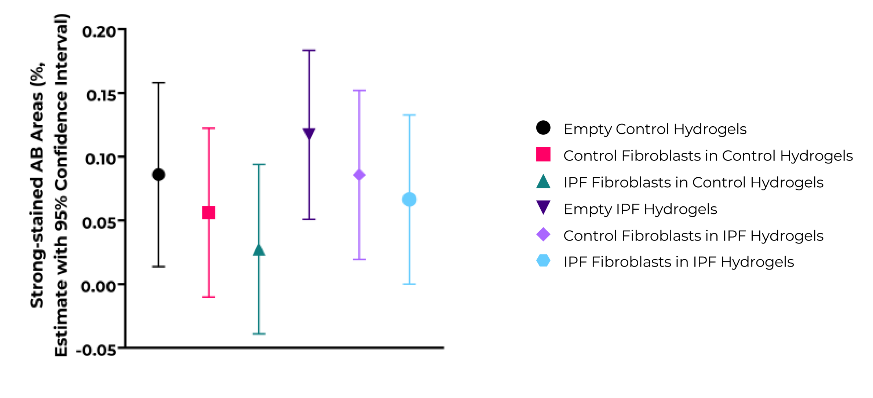

Supplementary Figure 23: Individual comparisons between the different groups of empty and fibroblast-encapsulated control and IPF hydrogels for the estimates of strongly stained Alcian blue (AB) area (%) for day 7 samples.

| Supplementary Table 18: Individual comparisons between the different groups of empty and fibroblast-encapsulated control and IPF hydrogels for strongly stained Alcian Blue area on day 14 samples | | | | | | | |
| --- | --- | --- | --- | --- | --- | --- | --- |
|  |  | Control Hydrogels | | | IPF Hydrogels | | |
|  |  | Empty | Control Fb | IPF Fb | Empty | Control Fb | IPF Fb |
| Control Hydrogels | Empty |  | 0.686 | 0.711 | 0.883 |  |  |
|  | Control Fb | 0.686 |  | 0.419 |  |  |  |
|  | IPF Fb | 0.711 | 0.419 |  |  |  |  |
| IPF Hydrogels | Empty | 0.883 |  |  |  | 0.253 | 0.358 |
|  | Control Fb |  |  |  | 0.253 |  | 0.818 |
|  | IPF Fb |  |  |  | 0.358 | 0.818 |  |
|  |  |  |  | Not significant | |  |  |
|  |  |  |  | Significant | |  |  |
|  |  |  |  | Comparison not applicable | |  |  |

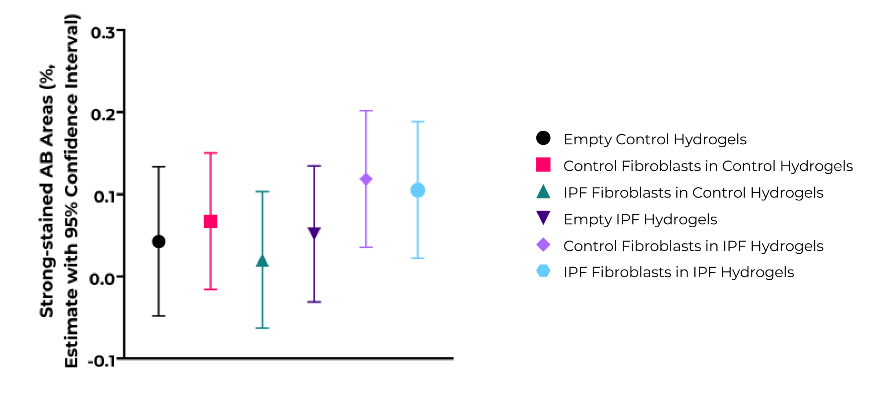

Supplementary Figure 24: Individual comparisons between the different groups of empty and fibroblast-encapsulated control and IPF hydrogels for the estimates of strongly stained Alcian blue (AB) area (%) for day 14 samples.

| Supplementary Table 19: Individual comparisons between the different groups of empty and fibroblast-encapsulated control and IPF hydrogels for high-density matrix on day 7 samples | | | | | | | |
| --- | --- | --- | --- | --- | --- | --- | --- |
|  |  | Control Hydrogels | | | IPF Hydrogels | | |
|  |  | Empty | Control Fb | IPF Fb | Empty | Control Fb | IPF Fb |
| Control Hydrogels | Empty |  | 0.400 | 0.412 | 0.630 |  |  |
|  | Control Fb | 0.400 |  | 0.109 |  |  |  |
|  | IPF Fb | 0.412 | 0.109 |  |  |  |  |
| IPF Hydrogels | Empty | 0.630 |  |  |  | 0.147 | 0.631 |
|  | Control Fb |  |  |  | 0.147 |  | 0.362 |
|  | IPF Fb |  |  |  | 0.631 | 0.362 |  |
|  |  |  |  | Not significant | |  |  |
|  |  |  |  | Significant | |  |  |
|  |  |  |  | Comparison not applicable | |  |  |

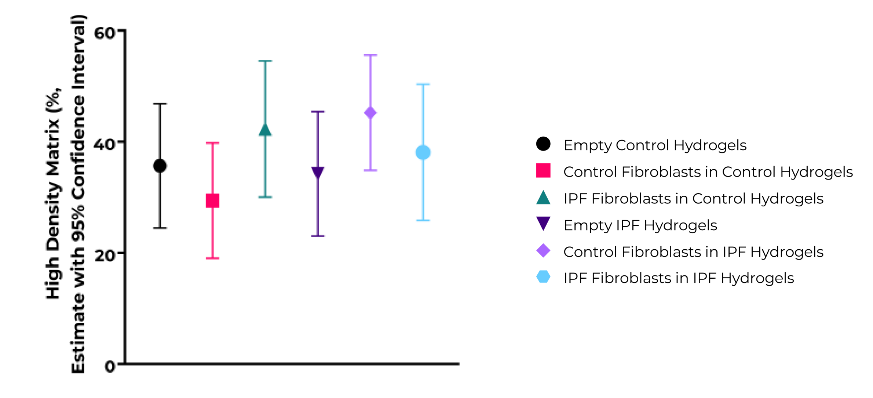

Supplementary Figure 25: Individual comparisons between the different groups of empty and fibroblast-encapsulated control and IPF hydrogels for the estimates of percentages of high-density matrix (%) for day 7 samples.

| Supplementary Table 20: Individual comparisons between the different groups of empty and fibroblast-encapsulated control and IPF hydrogels for high-density matrix on day 14 samples | | | | | | | |
| --- | --- | --- | --- | --- | --- | --- | --- |
|  |  | Control Hydrogels | | | IPF Hydrogels | | |
|  |  | Empty | Control Fb | IPF Fb | Empty | Control Fb | IPF Fb |
| Control Hydrogels | Empty |  | 0.512 | 0.056 | 0.029 |  |  |
|  | Control Fb | 0.512 |  | 0.160 |  |  |  |
|  | IPF Fb | 0.056 | 0.160 |  |  |  |  |
| IPF Hydrogels | Empty | 0.029 |  |  |  | 0.043 | 0.029 |
|  | Control Fb |  |  |  | 0.043 |  | 0.705 |
|  | IPF Fb |  |  |  | 0.029 | 0.705 |  |
|  |  |  |  | Not significant | |  |  |
|  |  |  |  | Significant | |  |  |
|  |  |  |  | Comparison not applicable | |  |  |

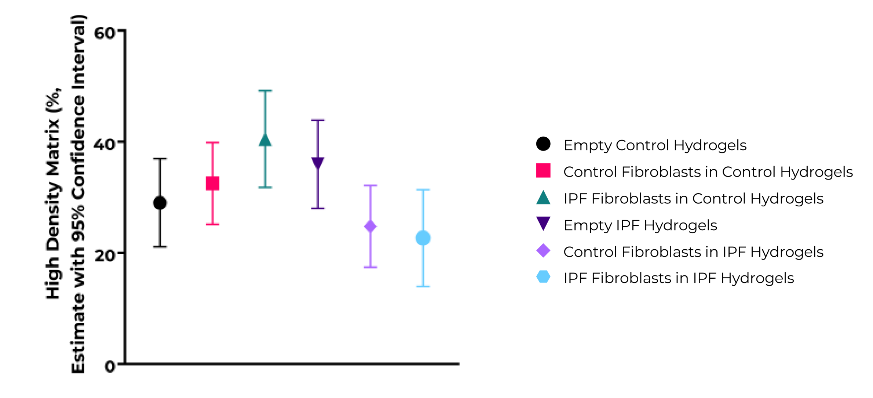

Supplementary Figure 26: Individual comparisons between the different groups of empty and fibroblast-encapsulated control and IPF hydrogels for the estimates of percentages of high-density matrix (%) for day 14 samples.

| Supplementary Table 21: Individual comparisons between the different groups of empty and fibroblast-encapsulated control and IPF hydrogels for fiber alignment on day 7 samples | | | | | | | |
| --- | --- | --- | --- | --- | --- | --- | --- |
|  |  | Control Hydrogels | | | IPF Hydrogels | | |
|  |  | Empty | Control Fb | IPF Fb | Empty | Control Fb | IPF Fb |
| Control Hydrogels | Empty |  | 0.266 | 0.401 | 0.572 |  |  |
|  | Control Fb | 0.266 |  | 0.844 |  |  |  |
|  | IPF Fb | 0.401 | 0.844 |  |  |  |  |
| IPF Hydrogels | Empty | 0.572 |  |  |  | 0.366 | 0.094 |
|  | Control Fb |  |  |  | 0.366 |  | 0.013 |
|  | IPF Fb |  |  |  | 0.094 | 0.013 |  |
|  |  |  |  | Not significant | |  |  |
|  |  |  |  | Significant | |  |  |
|  |  |  |  | Comparison not applicable | |  |  |

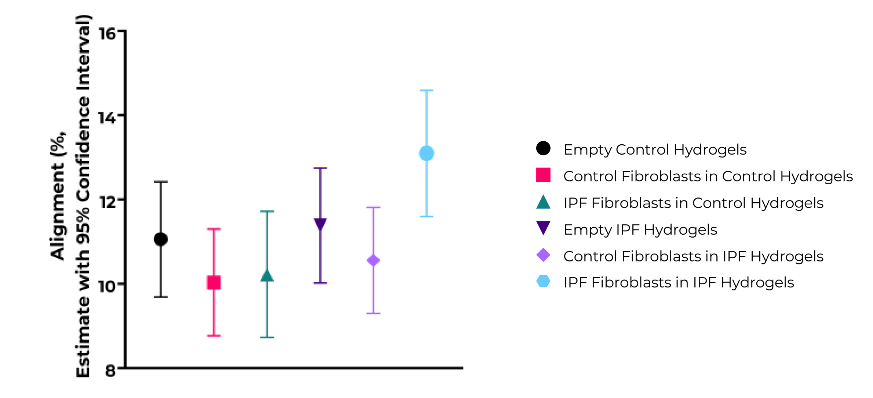

Supplementary Figure 27: Individual comparisons between the different groups of empty and fibroblast-encapsulated control and IPF hydrogels for the estimates of percentages of alignment (%) for day 7 samples.

| Supplementary Table 22: Individual comparisons between the different groups of empty and fibroblast-encapsulated control and IPF hydrogels for fiber alignment on day 14 samples | | | | | | | |
| --- | --- | --- | --- | --- | --- | --- | --- |
|  |  | Control Hydrogels | | | IPF Hydrogels | | |
|  |  | Empty | Control Fb | IPF Fb | Empty | Control Fb | IPF Fb |
| Control Hydrogels | Empty |  | 0.392 | 0.999 | 0.317 |  |  |
|  | Control Fb | 0.392 |  | 0.416 |  |  |  |
|  | IPF Fb | 0.999 | 0.416 |  |  |  |  |
| IPF Hydrogels | Empty | 0.317 |  |  |  | 0.094 | 0.019 |
|  | Control Fb |  |  |  | 0.094 |  | 0.335 |
|  | IPF Fb |  |  |  | 0.019 | 0.335 |  |
|  |  |  |  | Not significant | |  |  |
|  |  |  |  | Significant | |  |  |
|  |  |  |  | Comparison not applicable | |  |  |

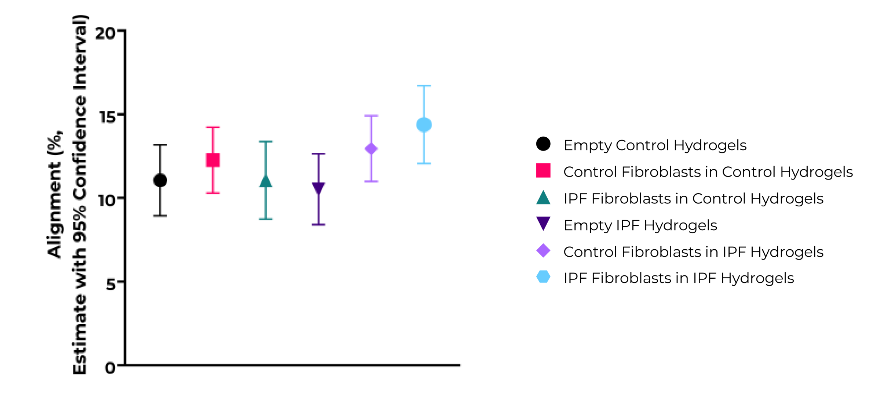

Supplementary Figure 28: Individual comparisons between the different groups of empty and fibroblast-encapsulated control and IPF hydrogels for the estimates of percentages of alignment (%) for day 14 samples.

| Supplementary Table 23: Individual comparisons between the different groups of empty and fibroblast-encapsulated control and IPF hydrogels for average fiber length on day 7 samples | | | | | | | |
| --- | --- | --- | --- | --- | --- | --- | --- |
|  |  | Control Hydrogels | | | IPF Hydrogels | | |
|  |  | Empty | Control Fb | IPF Fb | Empty | Control Fb | IPF Fb |
| Control Hydrogels | Empty |  | 0.311 | 0.689 | 0.004 |  |  |
|  | Control Fb | 0.311 |  | 0.175 |  |  |  |
|  | IPF Fb | 0.689 | 0.175 |  |  |  |  |
| IPF Hydrogels | Empty | 0.004 |  |  |  | 0.137 | 0.802 |
|  | Control Fb |  |  |  | 0.137 |  | 0.240 |
|  | IPF Fb |  |  |  | 0.802 | 0.240 |  |
|  |  |  |  | Not significant | |  |  |
|  |  |  |  | Significant | |  |  |
|  |  |  |  | Comparison not applicable | |  |  |

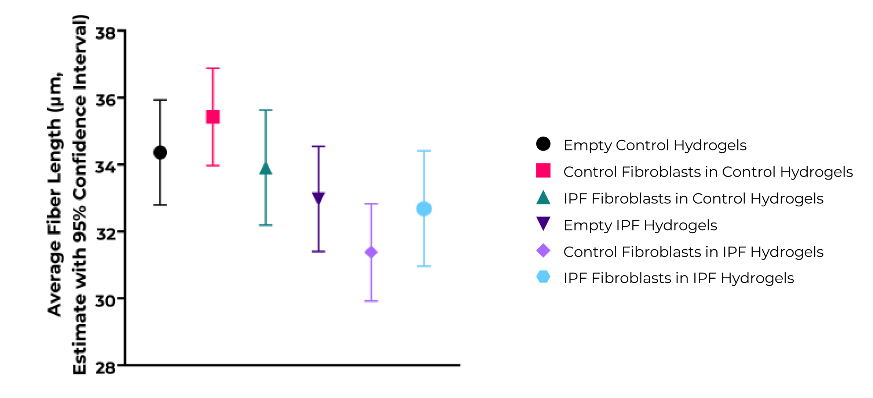

Supplementary Figure 29: Individual comparisons between the different groups of empty and fibroblast-encapsulated control and IPF hydrogels for the estimates of average fiber length (μm) for day 7 samples.

| Supplementary Table 24: Individual comparisons between the different groups of empty and fibroblast-encapsulated control and IPF hydrogels for average fiber length on day 14 samples | | | | | | | |
| --- | --- | --- | --- | --- | --- | --- | --- |
|  |  | Control Hydrogels | | | IPF Hydrogels | | |
|  |  | Empty | Control Fb | IPF Fb | Empty | Control Fb | IPF Fb |
| Control Hydrogels | Empty |  | 0.181 | 0.295 | 5.87 x 10⁻⁴ |  |  |
|  | Control Fb | 0.181 |  | 0.025 |  |  |  |
|  | IPF Fb | 0.295 | 0.025 |  |  |  |  |
| IPF Hydrogels | Empty | 5.87 x 10⁻⁴ |  |  |  | 0.263 | 0.123 |
|  | Control Fb |  |  |  | 0.263 |  | 0.571 |
|  | IPF Fb |  |  |  | 0.123 | 0.571 |  |
|  |  |  |  | Not significant | |  |  |
|  |  |  |  | Significant | |  |  |
|  |  |  |  | Comparison not applicable | |  |  |

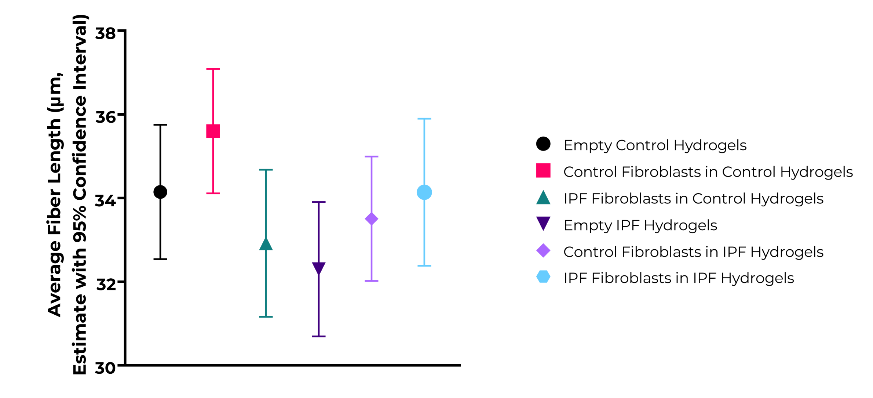

Supplementary Figure 30: Individual comparisons between the different groups of empty and fibroblast-encapsulated control and IPF hydrogels for the estimates of average fiber length (μm) for day 14 samples.

| Supplementary Table 25: Individual comparisons between the different groups of empty and fibroblast-encapsulated control and IPF hydrogels for number of endpoints on day 7 samples | | | | | | | |
| --- | --- | --- | --- | --- | --- | --- | --- |
|  |  | Control Hydrogels | | | IPF Hydrogels | | |
|  |  | Empty | Control Fb | IPF Fb | Empty | Control Fb | IPF Fb |
| Control Hydrogels | Empty |  | 0.259 | 0.155 | 1.68 x 10⁻⁴ |  |  |
|  | Control Fb | 0.259 |  | 0.033 |  |  |  |
|  | IPF Fb | 0.155 | 0.033 |  |  |  |  |
| IPF Hydrogels | Empty | 1.68 x 10⁻⁴ |  |  |  | 0.216 | 0.111 |
|  | Control Fb |  |  |  | 0.216 |  | 0.931 |
|  | IPF Fb |  |  |  | 0.111 | 0.931 |  |
|  |  |  |  | Not significant | |  |  |
|  |  |  |  | Significant | |  |  |
|  |  |  |  | Comparison not applicable | |  |  |

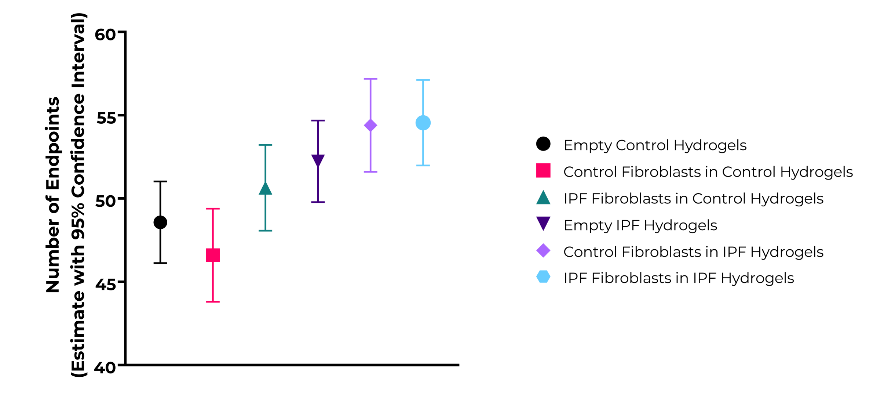

Supplementary Figure 31: Individual comparisons between the different groups of empty and fibroblast-encapsulated control and IPF hydrogels for the estimates of number of endpoints for day 7 samples.

| Supplementary Table 26: Individual comparisons between the different groups of empty and fibroblast-encapsulated control and IPF hydrogels for number of endpoints on day 14 samples | | | | | | | |
| --- | --- | --- | --- | --- | --- | --- | --- |
|  |  | Control Hydrogels | | | IPF Hydrogels | | |
|  |  | Empty | Control Fb | IPF Fb | Empty | Control Fb | IPF Fb |
| Control Hydrogels | Empty |  | 0.110 | 0.470 | 0.006 |  |  |
|  | Control Fb | 0.110 |  | 0.039 |  |  |  |
|  | IPF Fb | 0.470 | 0.039 |  |  |  |  |
| IPF Hydrogels | Empty | 0.006 |  |  |  | 0.790 | 0.260 |
|  | Control Fb |  |  |  | 0.790 |  | 0.505 |
|  | IPF Fb |  |  |  | 0.260 | 0.505 |  |
|  |  |  |  | Not significant | |  |  |
|  |  |  |  | Significant | |  |  |
|  |  |  |  | Comparison not applicable | |  |  |

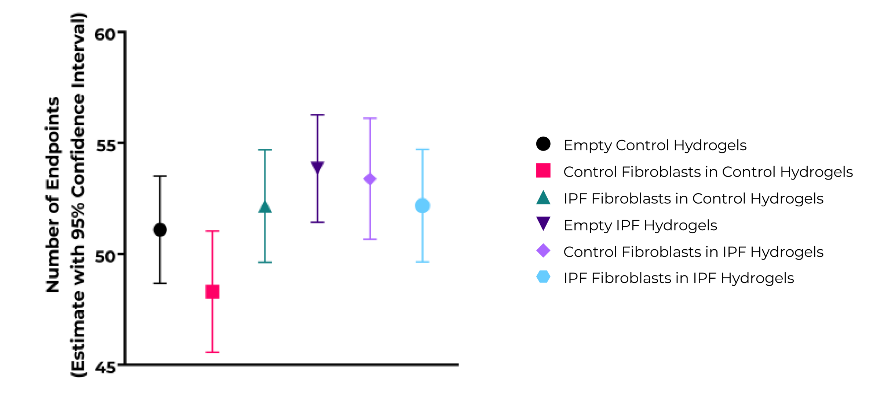

Supplementary Figure 32: Individual comparisons between the different groups of empty and fibroblast-encapsulated control and IPF hydrogels for the estimates of number of endpoints for day 14 samples.

| Supplementary Table 27: Individual comparisons between the different groups of empty and fibroblast-encapsulated control and IPF hydrogels for number of branchpoints on day 7 samples | | | | | | | |
| --- | --- | --- | --- | --- | --- | --- | --- |
|  |  | Control Hydrogels | | | IPF Hydrogels | | |
|  |  | Empty | Control Fb | IPF Fb | Empty | Control Fb | IPF Fb |
| Control Hydrogels | Empty |  | 0.719 | 0.895 | 0.007 |  |  |
|  | Control Fb | 0.719 |  | 0.633 |  |  |  |
|  | IPF Fb | 0.895 | 0.633 |  |  |  |  |
| IPF Hydrogels | Empty | 0.007 |  |  |  | 0.769 | 0.742 |
|  | Control Fb |  |  |  | 0.769 |  | 0.537 |
|  | IPF Fb |  |  |  | 0.742 | 0.537 |  |
|  |  |  |  | Not significant | |  |  |
|  |  |  |  | Significant | |  |  |
|  |  |  |  | Comparison not applicable | |  |  |

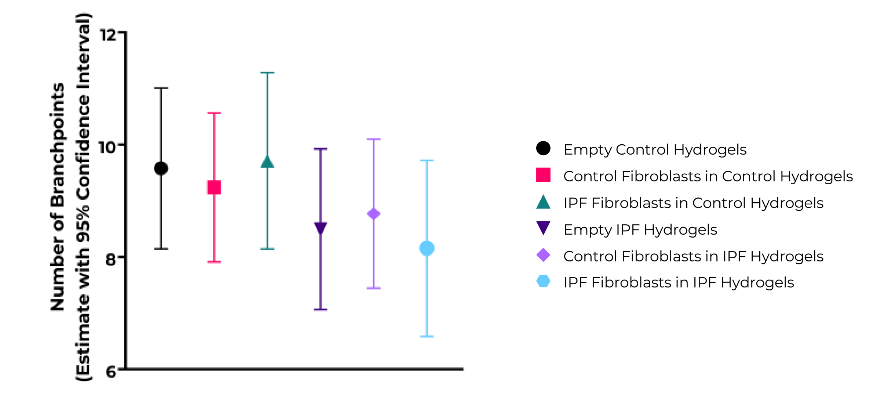

Supplementary Figure 33: Individual comparisons between the different groups of empty and fibroblast-encapsulated control and IPF hydrogels for the estimates of number of branchpoints for day 7 samples.

| Supplementary Table 28: Individual comparisons between the different groups of empty and fibroblast-encapsulated control and IPF hydrogels for number of branchpoints on day 14 samples | | | | | | | |
| --- | --- | --- | --- | --- | --- | --- | --- |
|  |  | Control Hydrogels | | | IPF Hydrogels | | |
|  |  | Empty | Control Fb | IPF Fb | Empty | Control Fb | IPF Fb |
| Control Hydrogels | Empty |  | 0.337 | 0.247 | 0.090 |  |  |
|  | Control Fb | 0.337 |  | 0.767 |  |  |  |
|  | IPF Fb | 0.247 | 0.767 |  |  |  |  |
| IPF Hydrogels | Empty | 0.090 |  |  |  | 0.071 | 0.148 |
|  | Control Fb |  |  |  | 0.071 |  | 0.798 |
|  | IPF Fb |  |  |  | 0.148 | 0.798 |  |
|  |  |  |  | Not significant | |  |  |
|  |  |  |  | Significant | |  |  |
|  |  |  |  | Comparison not applicable | |  |  |

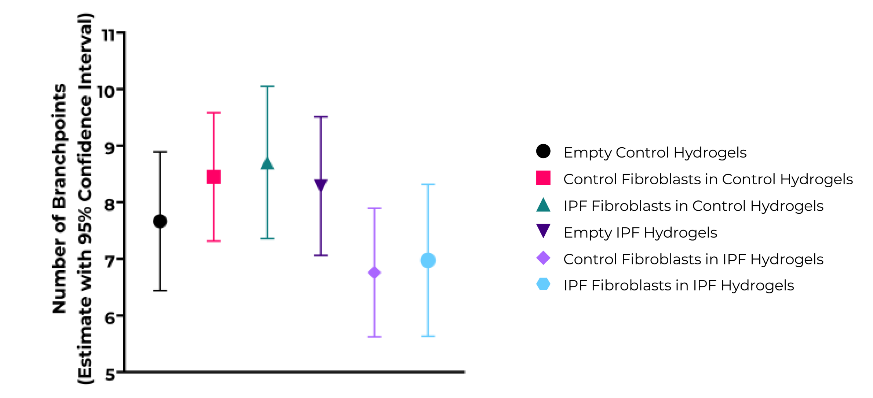

Supplementary Figure 34: Individual comparisons between the different groups of empty and fibroblast-encapsulated control and IPF hydrogels for the estimates of number of branchpoints for day 14 samples.

| Supplementary Table 29: Individual comparisons between the different groups of empty and fibroblast-encapsulated control and IPF hydrogels for low curvature window on day 7 samples | | | | | | | |
| --- | --- | --- | --- | --- | --- | --- | --- |
|  |  | Control Hydrogels | | | IPF Hydrogels | | |
|  |  | Empty | Control Fb | IPF Fb | Empty | Control Fb | IPF Fb |
| Control Hydrogels | Empty |  | 0.973 | 0.586 | 0.004 |  |  |
|  | Control Fb | 0.973 |  | 0.552 |  |  |  |
|  | IPF Fb | 0.586 | 0.552 |  |  |  |  |
| IPF Hydrogels | Empty | 0.004 |  |  |  | 0.043 | 0.871 |
|  | Control Fb |  |  |  | 0.043 |  | 0.074 |
|  | IPF Fb |  |  |  | 0.871 | 0.074 |  |
|  |  |  |  | Not significant | |  |  |
|  |  |  |  | Significant | |  |  |
|  |  |  |  | Comparison not applicable | |  |  |

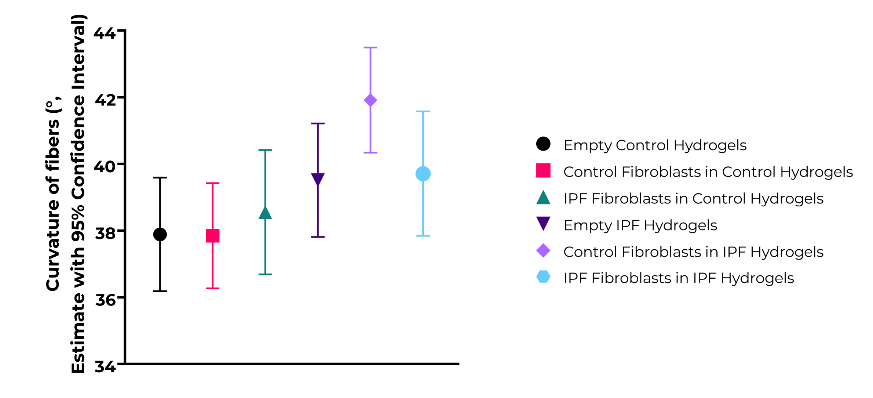

Supplementary Figure 35: Individual comparisons between the different groups of empty and fibroblast-encapsulated control and IPF hydrogels for the estimates of curvature of fibers in low curvature windows (°) for day 7 samples.

| Supplementary Table 30: Individual comparisons between the different groups of empty and fibroblast-encapsulated control and IPF hydrogels for low curvature window on day 14 samples | | | | | | | |
| --- | --- | --- | --- | --- | --- | --- | --- |
|  |  | Control Hydrogels | | | IPF Hydrogels | | |
|  |  | Empty | Control Fb | IPF Fb | Empty | Control Fb | IPF Fb |
| Control Hydrogels | Empty |  | 0.586 | 0.010 | 1.14 x 10⁻¹¹ |  |  |
|  | Control Fb | 0.586 |  | 0.027 |  |  |  |
|  | IPF Fb | 0.010 | 0.027 |  |  |  |  |
| IPF Hydrogels | Empty | 1.14 x 10⁻¹¹ |  |  |  | 0.018 | 0.009 |
|  | Control Fb |  |  |  | 0.018 |  | 0.593 |
|  | IPF Fb |  |  |  | 0.009 | 0.593 |  |
|  |  |  |  | Not significant | |  |  |
|  |  |  |  | Significant | |  |  |
|  |  |  |  | Comparison not applicable | |  |  |

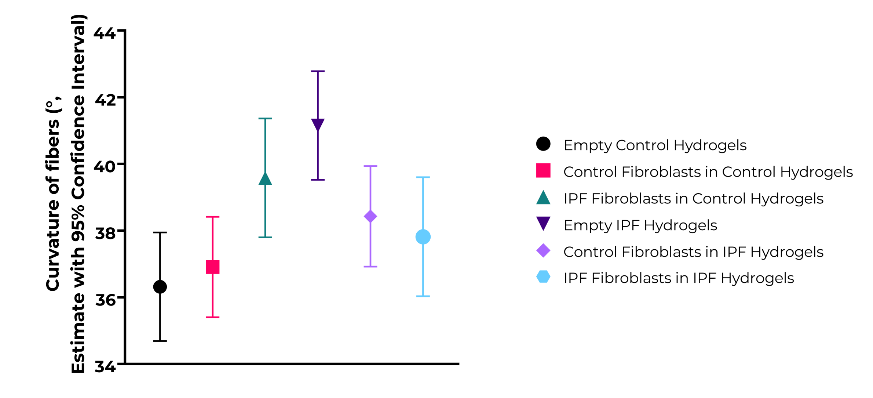

Supplementary Figure 36: Individual comparisons between the different groups of empty and fibroblast-encapsulated control and IPF hydrogels for the estimates of curvature of fibers in low curvature windows (°) for day 14 samples.

| Supplementary Table 31: Individual comparisons between the different groups of empty and fibroblast-encapsulated control and IPF hydrogels for high curvature window on day 7 samples | | | | | | | |
| --- | --- | --- | --- | --- | --- | --- | --- |
|  |  | Control Hydrogels | | | IPF Hydrogels | | |
|  |  | Empty | Control Fb | IPF Fb | Empty | Control Fb | IPF Fb |
| Control Hydrogels | Empty |  | 0.857 | 0.511 | 0.460 |  |  |
|  | Control Fb | 0.857 |  | 0.610 |  |  |  |
|  | IPF Fb | 0.511 | 0.610 |  |  |  |  |
| IPF Hydrogels | Empty | 0.460 |  |  |  | 0.345 | 0.171 |
|  | Control Fb |  |  |  | 0.345 |  | 0.025 |
|  | IPF Fb |  |  |  | 0.171 | 0.025 |  |
|  |  |  |  | Not significant | |  |  |
|  |  |  |  | Significant | |  |  |
|  |  |  |  | Comparison not applicable | |  |  |

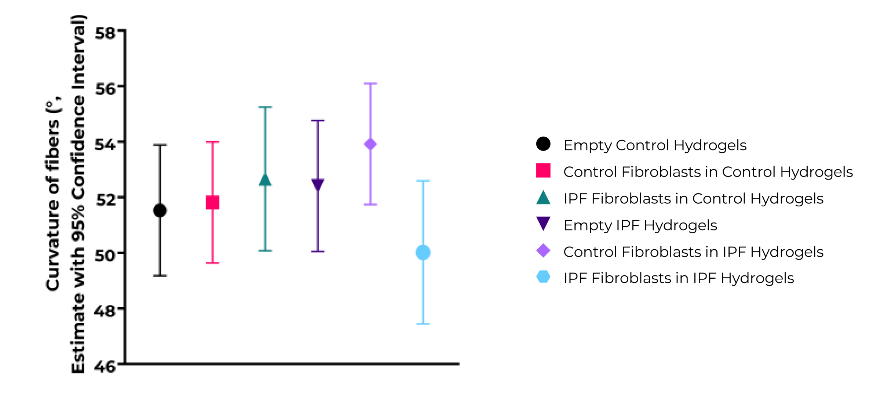

Supplementary Figure 37: Individual comparisons between the different groups of empty and fibroblast-encapsulated control and IPF hydrogels for the estimates of curvature of fibers in high curvature windows (°) for day 7 samples.

| Supplementary Table 32: Individual comparisons between the different groups of empty and fibroblast-encapsulated control and IPF hydrogels for high curvature window on day 14 samples | | | | | | | |
| --- | --- | --- | --- | --- | --- | --- | --- |
|  |  | Control Hydrogels | | | IPF Hydrogels | | |
|  |  | Empty | Control Fb | IPF Fb | Empty | Control Fb | IPF Fb |
| Control Hydrogels | Empty |  | 0.863 | 0.093 | 0.026 |  |  |
|  | Control Fb | 0.863 |  | 0.114 |  |  |  |
|  | IPF Fb | 0.093 | 0.114 |  |  |  |  |
| IPF Hydrogels | Empty | 0.026 |  |  |  | 0.780 | 0.058 |
|  | Control Fb |  |  |  | 0.780 |  | 0.766 |
|  | IPF Fb |  |  |  | 0.058 | 0.766 |  |
|  |  |  |  | Not significant | |  |  |
|  |  |  |  | Significant | |  |  |
|  |  |  |  | Comparison not applicable | |  |  |

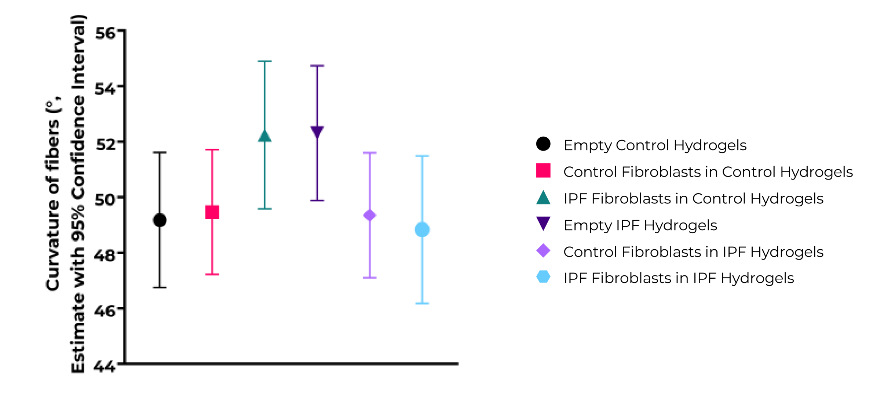

Supplementary Figure 38: Individual comparisons between the different groups of empty and fibroblast-encapsulated control and IPF hydrogels for the estimates of curvature of fibers in high curvature windows (°) for day 14 samples.

| Supplementary Table 33: Individual comparisons between the different groups of empty and fibroblast-encapsulated control and IPF hydrogels for stiffness on day 7 samples | | | | | | | |
| --- | --- | --- | --- | --- | --- | --- | --- |
|  |  | Control Hydrogels | | | IPF Hydrogels | | |
|  |  | Empty | Control Fb | IPF Fb | Empty | Control Fb | IPF Fb |
| Control Hydrogels | Empty |  | 0.497 | 0.846 | 4.93 x 10⁻²² |  |  |
|  | Control Fb | 0.497 |  | 0.639 |  |  |  |
|  | IPF Fb | 0.846 | 0.639 |  |  |  |  |
| IPF Hydrogels | Empty | 4.93 x 10⁻²² |  |  |  | 0.002 | 0.006 |
|  | Control Fb |  |  |  | 0.002 |  | 0.658 |
|  | IPF Fb |  |  |  | 0.006 | 0.658 |  |
|  |  |  |  | Not significant | |  |  |
|  |  |  |  | Significant | |  |  |
|  |  |  |  | Comparison not applicable | |  |  |

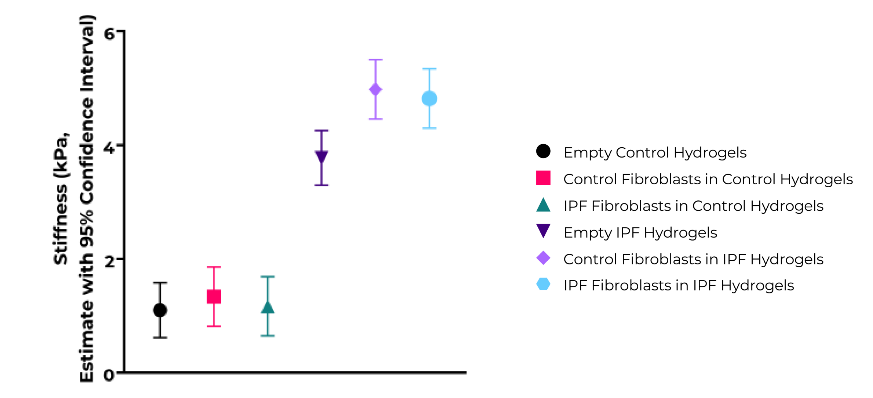

Supplementary Figure 39: Individual comparisons between the different groups of empty and fibroblast-encapsulated control and IPF hydrogels for the estimates of stiffness of hydrogels (kPa) for day 7 samples.

| Supplementary Table 34: Individual comparisons between the different groups of empty and fibroblast-encapsulated control and IPF hydrogels for stiffness on day 14 samples | | | | | | | |
| --- | --- | --- | --- | --- | --- | --- | --- |
|  |  | Control Hydrogels | | | IPF Hydrogels | | |
|  |  | Empty | Control Fb | IPF Fb | Empty | Control Fb | IPF Fb |
| Control Hydrogels | Empty |  | 0.958 | 0.653 | 1.15 x 10⁻³¹ |  |  |
|  | Control Fb | 0.958 |  | 0.629 |  |  |  |
|  | IPF Fb | 0.653 | 0.629 |  |  |  |  |
| IPF Hydrogels | Empty | 1.15 x 10⁻³¹ |  |  |  | 1.70 x 10⁻⁴ | 0.027 |
|  | Control Fb |  |  |  | 1.70 x 10⁻⁴ |  | 0.068 |
|  | IPF Fb |  |  |  | 0.027 | 0.068 |  |
|  |  |  |  | Not significant | |  |  |
|  |  |  |  | Significant | |  |  |
|  |  |  |  | Comparison not applicable | |  |  |

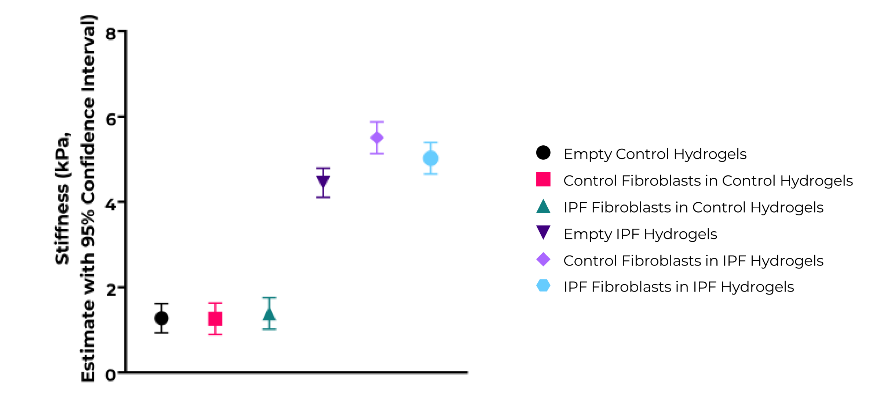

Supplementary Figure 40: Individual comparisons between the different groups of empty and fibroblast-encapsulated control and IPF hydrogels for the estimates of stiffness of hydrogels (kPa) for day 14 samples.

| Supplementary Table 35: Individual comparisons between the different groups of empty and fibroblast-encapsulated control and IPF hydrogels for time to reach 50% stress relaxation on day 7 samples | | | | | | | |
| --- | --- | --- | --- | --- | --- | --- | --- |
|  |  | Control Hydrogels | | | IPF Hydrogels | | |
|  |  | Empty | Control Fb | IPF Fb | Empty | Control Fb | IPF Fb |
| Control Hydrogels | Empty |  | 0.745 | 0.821 | 1.47 x 10⁻¹² |  |  |
|  | Control Fb | 0.745 |  | 0.924 |  |  |  |
|  | IPF Fb | 0.821 | 0.924 |  |  |  |  |
| IPF Hydrogels | Empty | 1.47 x 10⁻¹² |  |  |  | 0.573 | 0.691 |
|  | Control Fb |  |  |  | 0.573 |  | 0.358 |
|  | IPF Fb |  |  |  | 0.691 | 0.358 |  |
|  |  |  |  | Not significant | |  |  |
|  |  |  |  | Significant | |  |  |
|  |  |  |  | Comparison not applicable | |  |  |

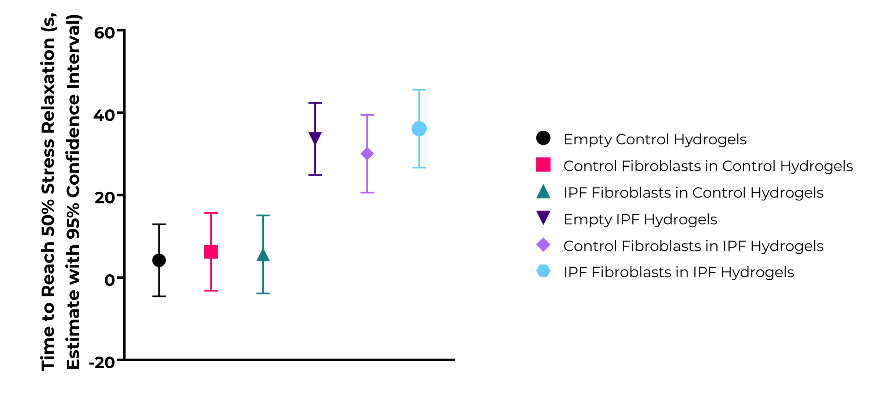

Supplementary Figure 41: Individual comparisons between the different groups of empty and fibroblast-encapsulated control and IPF hydrogels for the estimates of time to reach 50% stress relaxation (s) for day 7 samples.

| Supplementary Table 36: Individual comparisons between the different groups of empty and fibroblast-encapsulated control and IPF hydrogels for time to reach 50% stress relaxation on day 14 samples | | | | | | | |
| --- | --- | --- | --- | --- | --- | --- | --- |
|  |  | Control Hydrogels | | | IPF Hydrogels | | |
|  |  | Empty | Control Fb | IPF Fb | Empty | Control Fb | IPF Fb |
| Control Hydrogels | Empty |  | 0.719 | 0.515 | 5.82 x 10⁻¹⁴ |  |  |
|  | Control Fb | 0.719 |  | 0.778 |  |  |  |
|  | IPF Fb | 0.515 | 0.778 |  |  |  |  |
| IPF Hydrogels | Empty | 5.82 x 10⁻¹⁴ |  |  |  | 0.023 | 0.020 |
|  | Control Fb |  |  |  | 0.023 |  | 0.953 |
|  | IPF Fb |  |  |  | 0.020 | 0.953 |  |
|  |  |  |  | Not significant | |  |  |
|  |  |  |  | Significant | |  |  |
|  |  |  |  | Comparison not applicable | |  |  |

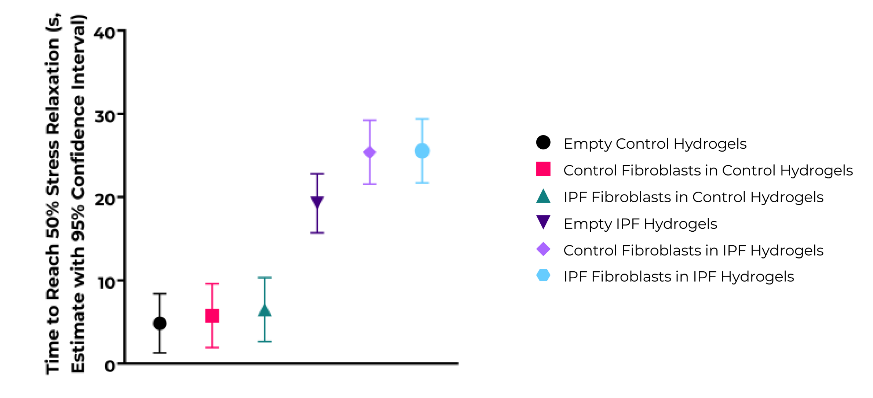

Supplementary Figure 42: Individual comparisons between the different groups of empty and fibroblast-encapsulated control and IPF hydrogels for the estimates of time to reach 50% stress relaxation (s) for day 14 samples.
